# *Drosophila grainyhead* gene and its neural stem cell-specific enhancers show epigenetic synchrony in the cells of the central nervous system

**DOI:** 10.1101/2025.03.20.644476

**Authors:** Rashmi Sipani, Yamini Rawal, Jiban Barman, Prakeerthi Abburi, Vishakha Kurlawala, Rohit Joshi

## Abstract

Enhancers are the epicentres of tissue-specific gene regulation. In this study, we have used the central nervous system (CNS) specific expression of the *Drosophila grainyhead* (*grh*) gene to make a case for deleting the enhancers in a sensitised background of other enhancer deletion, to functionally validate their role in tissue-specific gene regulation. We identified novel enhancers for *grh* and subsequently deleted two of them, to establish their collective importance in regulating *grh* expression in CNS. This showed that *grh* relies on multiple enhancers for its robust expression in neural stem cells (NSCs), with different combinations of enhancers playing a critical role in regulating its expression in various subset of these cells. We also found that these enhancers and the *grh* gene show epigenetic synchrony across the three cell types (NSCs, intermediate progenitors and neurons) of the developing CNS; and *grh* is not transcribed in intermediate progenitor cells, which inherits the Grh protein from the NSCs. We propose that this could be a general mechanism for regulating the expression of cell fate determinant protein in intermediate progenitor cells. Lastly, our results underline that enhancer redundancy results in phenotypic robustness in *grh* gene expression, which seems to be a consequence of the cumulative activity of multiple enhancers.

## Introduction

The metazoan gene transcription is highly complex and relies on multiple enhancers (Cis-Regulatory Elements-**CREs**) to fine-tune spatial and temporal dynamics of the gene expression. The *lacZ* reporter assays (used for testing the enhancer activity of genomic fragments by fusing them to a reporter protein like β-galactosidase) have been standard for identifying new CREs. However, besides enhancer deletion, there is no easy way to establish whether a CRE is required for gene expression in a particular tissue. With the advent of CRISPR-Cas9 technology [1–3], it is now possible to precisely delete CREs and address their role in tissue-specific gene regulation and development. Such functional validation will give credibility to the associated biochemical, genetic and epigenetic experiments and provide physiologically relevant insights into gene regulation.

*Drosophila* offers sophisticated genetic tools like CRISPR-Cas9 [4, 5], which enables precise deletion of the enhancer, and Targeted DamID (**TaDa**) [6], which allows probing the chromatin status of the gene/enhancer in a tissue-specific manner [7]. However, despite these advances, the in-vivo cell-specific chromatin status of a functionally validated tissue-specific enhancer has not been probed. Similarly, how the epigenetic status of a gene correlates with cell-specific enhancers and how it changes across different cell types of developing central nervous system (**CNS**) also remains unexplored. In this work, we tried to address these questions by studying neural stem cell (**NSC**) specific expression of *Drosophila grainyhead* gene.

*Drosophila grainyhead* (***grh***) and its vertebrate orthologs (Grainyhead-like, **Grhl-1-3)** code for basic helix-loop-helix transcription factor (**TF**) [8, 9] [10] [11] [12–14] which function as a master regulators for maintenance and homeostasis of epithelial cells [12–17]. In *Drosophila* CNS, which comprises of optic lobe (OL), central brain (CB) and ventral nerve cord (VNC) (Fig 1A), Grh plays a role in the generation of neuronal diversity by functioning as a temporal series transcription factor (**tTF**) both in Type I NSCs of VNCs and intermediate progenitors of Type II NSCs in CB (Fig 1A) [21–25]. Type I NSCs, which account for 95% of neural stem cells, divide asymmetrically to self-renew and give rise to intermediate progenitor (or Ganglion Mother Cells-**GMCs**). The GMC then divides symmetrically to give rise to a pair of neurons or glia. The Type-II NSCs also divide asymmetrically to self-renew and give rise to intermediate progenitors (called INPs); these INPs then divide like Type-I NSCs, thereby giving rise to many more progeny in a lineage. There are only 16 Type-II NSCs in *Drosophila,* which are very similar to mammalian cortical NSCs. Grh is expressed in the Type I NSCs and their GMCs as early as stage 15 of embryogenesis [8]. In Type II NSCs and their Intermediate Progenitor Cells (**INPs**), Grh is expressed during the larval stages [22–25]. Grh has also been shown to play a role in the NSC proliferation [13, 26, 27] and their Hox-dependent apoptosis in the abdominal and terminal (A3-A10) segments of larval CNS [26, 28–32]. More recently, *Drosophila* Grh was also shown to function as a pioneer TF regulating the chromatin accessibility of its target genes [13, 14, 18–20]. However, despite their importance in neural and non-neural tissues, little is known about the mechanisms regulating the expression of Grh and its vertebrate orthologs [33–35]. In vertebrates, a recent study identified and deleted a highly conserved 2.7 Kb enhancer (*mm1286*) that drives the expression of *grhl2* in craniofacial primordia during embryogenesis. However, the deletion of this enhancer showed no defect in craniofacial development or palatal closure [34]. Similarly, in *Drosophila*, only two genomic regions are known to regulate the *grh* expression in NSCs of CNS, while none has been identified for epithelial tissue. Neither of the two enhancers have been deleted to functionally test for their role in regulating *grh* expression in NSCs.

**Fig 1.**
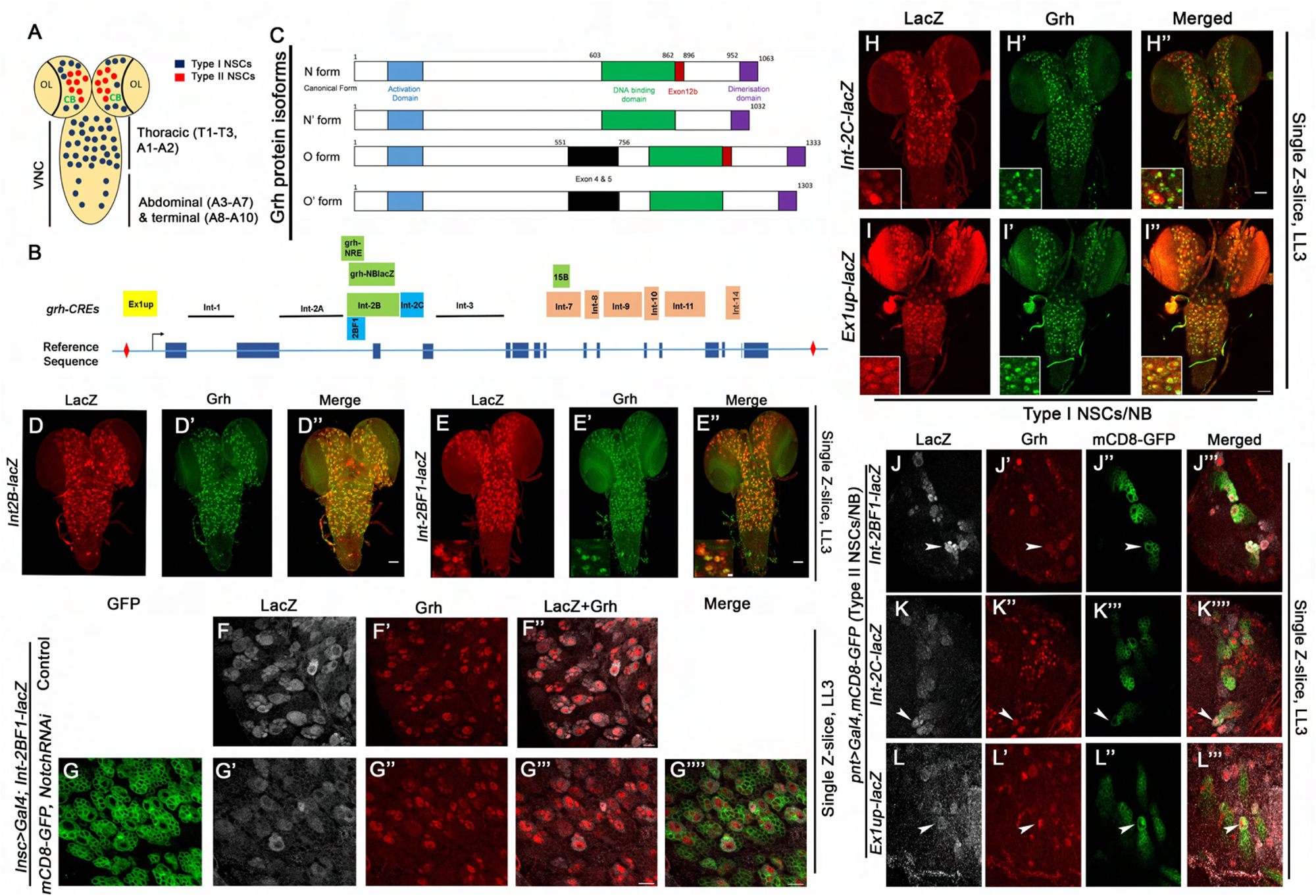
Expression analysis of *grh* enhancers. (A) Schematic of larval CNS showing optic lobes (OL), central brain (CB) and ventral nerve cord (VNC). (B) Schematic of the genomic region of *grh* (exons are denoted by filled blue boxes, and blue lines denote the introns). All the genomics regions tested by reporter lacZ assay are shown at the top of the reference sequence. Black lines denote the regions which showed no expression in larval NSCs, and coloured boxes show regions that exhibited expression in late L3 stage NSCs. The known *grh* enhancers like 4 Kb *grh-NB-lacZ* or *Int-2B-lacZ* (covering 1.3 Kb *grh-NRE*) and 0.8 Kb *15B* are shown in green coloured boxes. One yellow and two cyan colored boxes show the positions of 2.7 Kb *Ex1up,* 1.2 Kb *Int-2BF1*, and 1.5 Kb *Int-2C*, respectively. Red diamonds and arrow indicate the approximate position of Type I insulator element and transcription start site.(C) Schematic for Grh N, N’, O and O’ isoforms (adapted from Uv et al)[13]. (D-E) 1.2 Kb subfragment (*Int-2BF1*) (E) recapitulates the activity of the 4 Kb enhancer *(Int-2B*) (D) in all the Grh expressing NBs. (F-G) In the Type I NSCs of the thoracic region, the *Int-2BF1-lacZ* is downregulated in response to Notch knockdown, but the levels of Grh protein are unaffected. (H-I) Novel CREs *Int-2C* and *Ex1up-lacZ* show activity in all the Grh-expressing NSCs at the late L3 stage. (J-L) *Int-2BF1, Int-2C* and *Ex1up-lacZ* are also expressed in Type-II NSCs (marked by *Pnt-GAL4>UAS-mCD8GFP*). NSCs (NBs) are marked with Deadpan (Dpn) in the late L3 stages. Notch knockdown in panels F-G is from the early embryonic stages and VNCs are analysed in late L3 (LL3) stage. White arrowheads indicate Type II NSCs. Scale bars are 50µm (for D, E, H and I) and 10µm (for F-G).

In this study, we identified seven new potential CREs that may regulate Grh expression in NSCs and wing primordium of *Drosophila*. Subsequently, we used CRISPR-Cas9 and deleted two of these enhancers to test their role in regulating the NSC-specific expression of the *grh* gene. The homozygotes for the single enhancer deletions were viable as adults and did not affect Grh expression in larval NSCs. But in the case where both the copies of the two enhancers were deleted (trans-heterozygotes for the enhancer double deletion-*grh^DD^* and *grh* deficiency-*grh^DD^/grh^Df^*), we observed a significant decrease in the expression of the Grh from a majority of NSCs and a complete loss of Grh from the remaining minority of NSCs. This also resulted in proliferation defect and a block of apoptosis of NSCs while leaving the expression of Grh in the non-neural tissue (wing disc) unaffected. These results suggest that the robustness of *grh* expression in CNS relies on multiple enhancers, and different combinations of these enhancers are required to critically regulate its expression in various subsets of cells. Our double deletion experiments also suggest that two enhancers deleted here are collectively important for regulating *grh* expression in developing CNS.

The epigenetic status of the deleted enhancers and *grh* gene showed synchrony across the three cell types of CNS, with *grh* expression being differentially regulated in NSCs by polycomb protein (Pc), and switched off in neurons by HP1a (Heterochromatin Protein 1a) and repressive Histone-H1. We also found that Grh is not transcribed in GMCs, which inherited Grh protein from NSCs; which could be a general strategy for regulating the expression of cell fate determinant protein in intermediate progenitor cells. Collectively, we report functional identification of one subset of enhancers important for regulating the *grh* expression in the NSCs of the CNS. Our deletion experiments underline the fact that enhancer redundancy results in the phenotypic robustness in *grh* gene expression, which seems to be a consequence of the cumulative activity of multiple enhancers.

## Results

### Expression analysis of *grh* enhancers

Only two enhancers are suggested to regulate *grh* expression in NSCs of the CNS of *Drosophila*; a 4 Kb genomic region located in the second intron [12, 36] and a 1.5 kb region in the seventh intron of the *grh* gene (indicated by the green color in Fig 1B) [37]. The 4 Kb region (called *grh-NB* in an earlier study [12, 36]) also covers a previously characterised 1.3 Kb *grh-NRE* enhancer [12, 36] and expressed strongly in larval NSCs (Fig 1B). We called this 4 Kb enhancer *Int-2B* owing to its location in the second intron (Fig 1B and S1A).

To identify the minimal enhancer region within the 4Kb CRE of *grh*, we subfragmented this region based on the sequence conservation. A 1.2 Kb sub-fragment of *Int-2B* (*Int-2BF1,* Fig 1B) was found to recapitulate the expression of the entire 4 Kb region, while the other two fragments (*Int-2BF2* and *Int-2BF3*) did not express in larval CNS. This 1.2 Kb region was further narrowed to 150 bps, which had to be trimerised to show significant activity in larval NSCs (Fig S1 and Sup Data). Since Grh protein and *grh-NRE* are known to be sensitive to Notch signalling in Type-II NSCs of larval CNS [36], we tested if the same happened for *Int-2BF1* in Type-I NSCs. We observed that though *Int-2BF1-lacZ* showed partial downregulation in NSCs, only a few cells showed Grh downregulation (Fig 1F and 1G’), suggesting that Grh expression in Type-I NSCs is largely insensitive to Notch signalling.

Next, based on sequence conservation across multiple *Drosophila* species we screened 11 non-coding genomic regions bracketed within two class I genomic insulator elements (indicated by red diamonds in Fig 1B) for *grh* gene [38] (detailed in sup Data). We identified seven new CREs (*Ex1up, Int-2C, Int-8, 9, 10, 11, 14* using reporter-lacZ assay) which expressed in the larval NSCs of the CNS and only one CRE (*Ex1up*) that expressed in non-neural wing disc tissue (Fig 1F-1L, Fig S2, Fig S3, expression also summarised in Table I).

In contrast to most NSCs in the central brain (CB) and thoracic region of the VNC, the NSCs in the A8-A10 segments (terminal region Fig 1A) do not express Grh in early larval stages and showed a temporal increase in Grh expression and Notch activity from mid-larval stages, resulting in their apoptosis [31]. Using this increase in Grh expression as a proxy for temporal dynamics of Grh, we found that five out of a total of nine CREs could successfully recapitulate Grh expression and dynamics in A8-A10 NSCs before they died in the mid-L3 stage (Table I). This included three of the new CREs (*Ex1up, Int-2C,* and *Int-14*) (Fig S4D-S4I and Table I) and two already known CREs (*Int-2BF1* and *Int-7*, Fig S4A-S4C, Table I).

Based on the uniformity, temporal dynamics and strength of their expression in larval NSCs, the enhancers (both novel and known [in bold]) were divided into three categories. The first category included the ones which showed strong and uniform expression in the majority of the NSCs (*Ex1up, **Int-2BF1**, Int-2C, Int-14*-highlighted in green in Table I) (Fig 1E, 1H-1I, Fig S2E and S2J). The second category included the CREs, which were very weak in their expression in NSCs (*Int-8, Int-9* and *Int-11*-highlighted in pink in Table I) (Fig S2B-S2D and S2G-S2I). The third category included the remaining two enhancers (***Int-7***, *Int-10*-highlighted in cyan in Table I), of which *Int-7* showed strong but variable expression in the NSCs, and *Int-10* showed strong expression in Type II NSCs and very limited and weak expression in Type I NSCs (detailed in Sup Data and Fig S2A and S2F, and S2K-S2L). This suggested that *Int-10* is an important CRE for the regulation of *grh* in Type II NSCs. Thereafter, we compared the expression of our *enhancer-lacZ* lines with the enhancer expression dataset available for *grh* gene in the *Drosophila* FlyLight database, which curates CNS-specific expression analysis of enhancers for select *Drosophila* genes. Our enhancer expression largely matched those of FlyLight lines (Sup Data).

Collectively, we have identified seven new CREs for the *grh* gene (*Ex1up, Int-2C, Int-8, 9, 10, 11, 14*), which, along with the existing two enhancers (*Int-2BF1* and *Int-7*), may play a role in regulating the expression of Grh in neural and non-neural tissues. Amongst the eleven genomic regions screened, we observed only one enhancer (*Ex1up*) expressed in epithelial wing primordium and one (*Int-10*) showing expression primarily in Type II NSCs. We also found that unlike Type II NSCs, Grh expression in Type I NSCs is insensitive to Notch signalling.

### Grh expression in NSCs is unaffected in individual deletion for *Int-2BF1* and *Int-2C*

Next, we focused on three enhancers from the first category (*Ex1up-2.7kb, **Int-2BF1-1.1kb*** and *Int-2C-1.5kb*) owing to their relative proximity and strong expression (Fig 2G). Using the double guide CRISPR-Cas9 approach [4] we could generate stable single deletions for *Int-2BF1* and *Int-2C* (*ΔInt-2BF1* and *ΔInt-2C*) (Fig 2A and 2B and Table II). However, both these single deletions were homozygous viable and fertile as adults and did not show any significant change in Grh expression in NSCs (Fig 2C-2F and C’’’ and E’’’).

**Fig 2.**
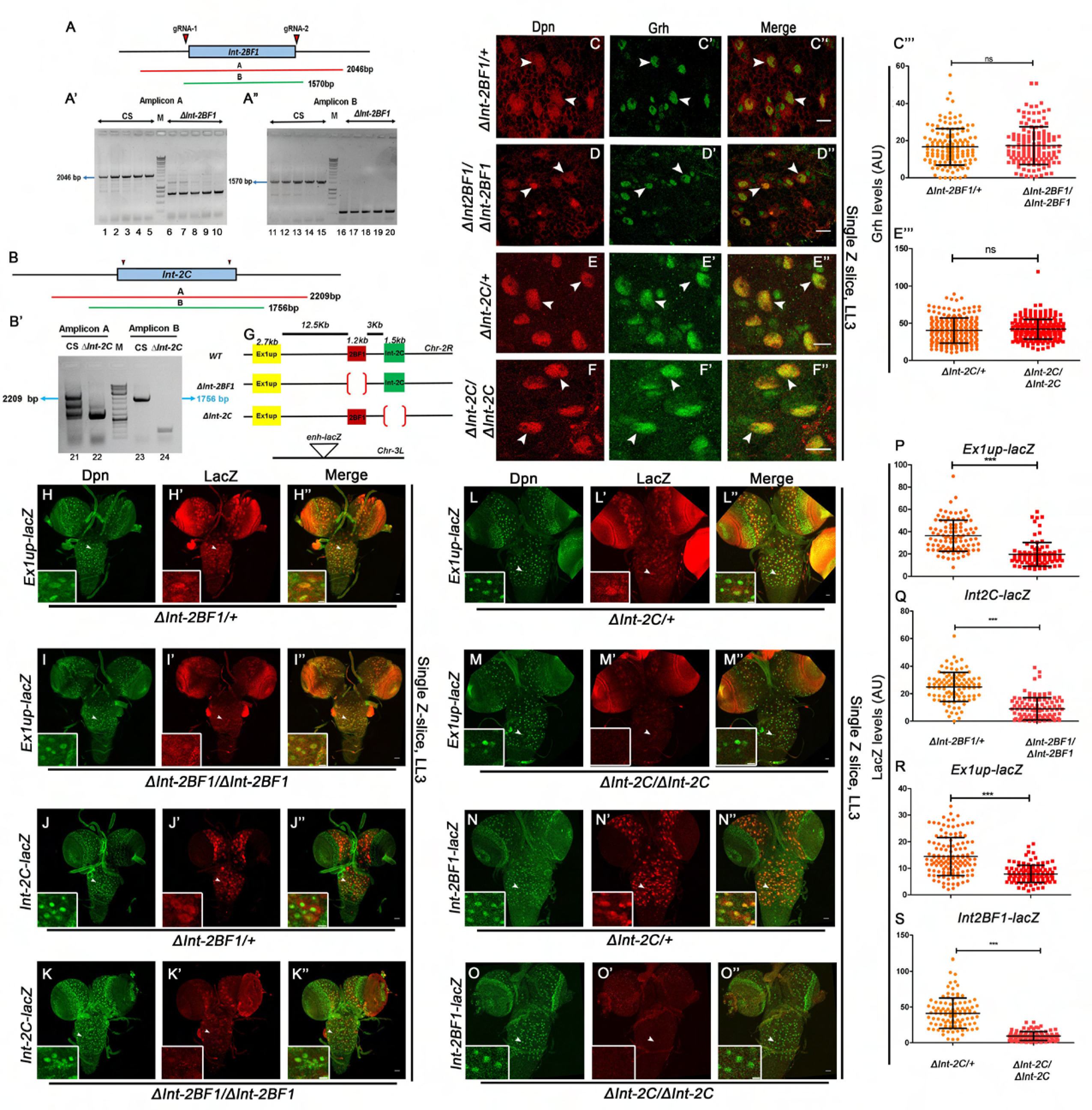
*grh* enhancers *Int-2BF1* and *Int-2C* show activity interaction. (A-B) Shows the approximate positions of the guide RNAs and the size of the amplicons used to screen the deletions of *Int-2BF1* (A) and *Int-2C* (B). Red and green lines show the extent of amplicons. (A’-B’) Shows the agarose gels confirming *Int-2BF1* (lanes 6 to 10 and 16-20) and *Int-2C* deletions (lanes 22 and 24) in the homozygous larvae. Canton-S genomic is used as a wild-type control. (C-F) Shows that the Grh expression is unaffected in the NSCs of heterozygous and homozygous deletions in the late L3 (LL3) stage deletions of *Int-2BF1* (C vs D) and *Int-2C* (E vs F). (C’’’ and E’’’) Show the quantitation graphs for Grh levels in heterozygous vs homozygous single deletions. (G) Schematic showing the approximate positions and sizes of *grh* enhancers *Ex1up* (yellow box)*, Int-2BF1* (maroon box)*, and Int-2C* (green box) on Chr-2R and site of insertion of different *enhancer-lacZ* construct on Chr-3L. (H-K) Compared to heterozygotes the homozygotes for *Int-2BF1* deletion (*ΔInt-2BF1*) show a significant reduction in the activity of *Ex1up-lacZ* (H vs I) and *Int-2C-lacZ* (J vs K) in late L3 (LL3) stage NSCs compared to the heterozygotes for the deletion. (L-O) Homozygotes for *Int-2C* deletion (*ΔInt-2C*) show a significant reduction in the expression of *Ex1up-lacZ* (L vs M) and *Int-2BF1-lacZ* (N vs O) in late L3 stage NSCs compared to the heterozygotes for the deletion. (P-S) Shows graphs for LacZ levels in heterozygous vs homozygous single deletions. White arrowheads indicate the NSCs (NBs) marked by Deadpan (Dpn) in panels C-F and approximate position of the high magnification insets in panels H-O. Scale bars are 20 µm (for C-F), 30 µm (H-O) and 10 µm for the insets.

The Grh have four isoforms, two neural (GrhO and O’) and two non-neural (GrhN and N’) (Fig 1C). The CNS-specific O-isoforms have two additional exons (exon-4 and 5), which code for 205 residues [13] (Fig-1C). The polyclonal Grh antibody used by us identified all the isoforms. Since *Int-2BF1* and *Int-2C* deletions showed no decrease in the levels of the Grh in NSCs, we considered the possibility that in addition to O-isoforms, CNS may be expressing N-isoforms as well, and decided to check single deletions for the expression of GrhO isoforms specifically. For this, we generated a GrhO-specific polyclonal antibody against 205 residues of exon-4 and 5 (Fig-1C). We found that the Grh Exon 4+5 antibody exclusively stained NSCs and GMCs in the CNS (inset Fig-S5A’) but not the cells in the eye (Fig S5C’ vs S5B’) and wing imaginal discs (Fig S5D and S5E), establishing that this antibody was specific for the GrhO isoform. However, we did not find any change in the levels of the Grh in the case of homozygotes for the single deletions even with O-isoform specific antibody (Fig-S5F-S5I). These observations collectively suggested that single enhancer deletions of *Int-2BF1* or *Int-2C* did not affect NSC-specific Grh expression.

CREs are often known to function redundantly to control the expression of a given gene. Therefore, upon deleting a particular CRE, the other CREs may increase their activity [39, 40]. To test this, we checked if the deletion of the *Int-2BF1* or *Int-2C* led to increased activity of the other CREs (Fig 2G). This was done by checking the expression of *enh-lacZ* lines in the background of single enhancer deletions (Fig 2G). The *enh-lacZ* lines were inserted at Chr-3L, a genomic location different from endogenous *grh* gene located on Chr-2R. To this end, we monitored the activity of *Ex1up-lacZ* and *Int-2C-lacZ* in the background of the deletion for *Int-2BF1 (ΔInt-2BF1)* (Fig 2H-2K) and *Ex1up-lacZ* and *Int-2BF1-lacZ* in the background of the deletion for *Int-2C (ΔInt-2C)* (Fig2L-2O). Contrary to our expectation, we observed that the activity of *Ex1up-lacZ* (Fig 2H-2I) and *Int-2C-lacZ* (Fig 2J-2K) was significantly downregulated in the case of *Int-2BF1* deletion homozygotes *(ΔInt-2BF1/ΔInt-2BF1)*. Similarly, the activity of *Ex1up-lacZ* (Fig 2L-2M) and *Int-2BF1-lacZ* (Fig 2N-2O) was also significantly downregulated in the case of *Int-2C* deletion homozygotes *(ΔInt-2C/ΔInt-2C)* (Fig 2P-2S). These results suggested that the three enhancers may be physically interacting with each other, which may be important for the individual enhancer activity and crucial for overall gene expression. Since *Int-2BF1* and *Int-2C* CREs are located close by in the second intron (approximately 3 Kb apart, Fig 2G), and the effect of *Int-2C* deletion on the expression of *Int-2BF1-lacZ* (on 3L) was most pronounced (Fig-2O’), we considered deleting these two CREs together.

### Double deletion of *Int-2BF1* and *Int-2C* reduces Grh expression in NSCs

To test the collaborative functioning of *Int-2BF1* and *Int-2C,* we made a double deletion for the two CREs. The double deletion was referred to as *ΔInt-2BF1-ΔInt-2C’* (*grh^DD^)* since the extent of *Int-2C* deletion here differed from the original *Int-2C* deletion (Table II). The double deletion was homozygous lethal as embryos, but the trans-heterozygotes of double deletion and *grh* deficiency (*grh^DD^/grh^Df^*) survived till the adult stage, indicating the existence of a third site modifier in the double deletion chromosome, which is a non-specific mutation outside *grh* locus. The *grh^DD^/grh^Df^*larval CNS showed a significant reduction in the expression of Grh from a majority of NSCs (Fig 3C vs 3C’ and white arrowheads in Fig 3E’ vs 3F’), with complete loss of Grh from a small subset of NSCs (yellow arrowheads, Fig 3F’ and 3G), while the expression of Grh in the cells of wing discs was unaltered (Fig 3D vs 3D’). This suggested *Int-2BF1* and *Int-2C* are important for regulating the Grh expression in the majority of the NSCs of the CNS and critical in a small minority of NSCs which completely lose the Grh.

**Fig 3.**
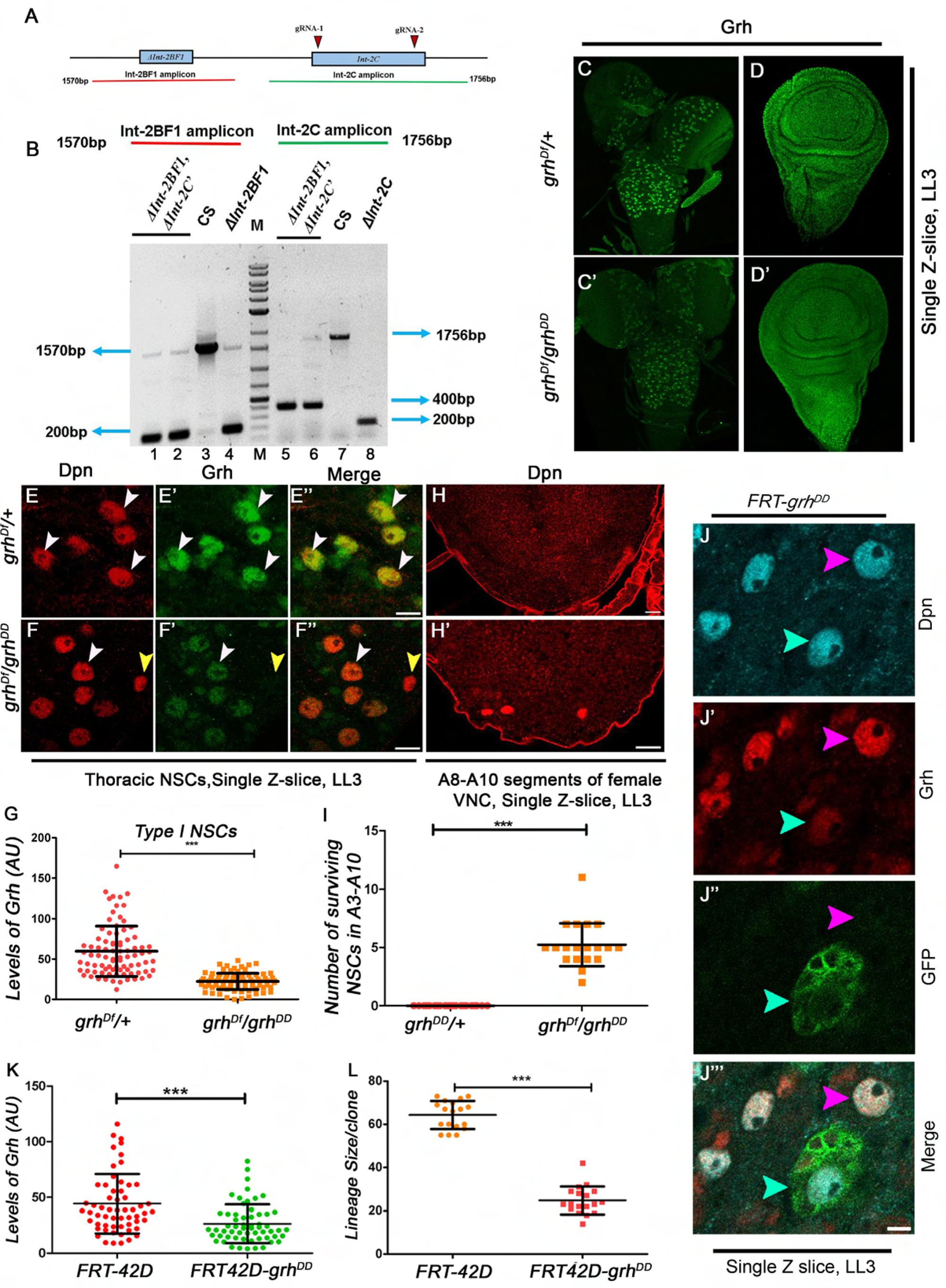
Double deletion of *Int-2BF1* and *Int-2C* enhancers (*grh^DD^*) compromises Grh expression in NSCs but not in the wing primordium. (A) Shows the approximate positions of the guide RNA sequences. Red and green lines show the extent of amplicons used for screening *Int-2BF1* and *Int-2C* deletions, respectively. *Int-2C* was deleted in the background of *Int-2BF1* deletion (*ΔInt-2BF1*). (B) Agarose gel confirming *Int-2BF1* (lanes 1 to 4) and *Int-2C* deletions (lanes 8 to 9) in the heterozygotes for the double deletion larvae (*ΔInt-2BF1, ΔInt-2C’/+* or *grh^DD^*). Canton-S genomic is used as wild-type control; the extent of *Int-2C* deletion is different from the one shown in Fig 2 and is referred to as *ΔInt-2C’*. (C and E-F) Compares the Grh expression in whole CNS (C vs C’) and thoracic NSCs of heterozygous *(grh^Df^/+)* (E-E’’) and heteroallelic combination of *grh* double deletion and deficiency (*grh^Df^/grh^DD^*) (F-F’’) in the late L3 (LL3) stage. The *grh^Df^/grh^DD^* shows a significant decrease in Grh expression in the NSCs (white arrowheads), with some NSCs showing a complete loss of Grh (yellow arrowheads), while the wing discs (D-D’) do not show any change in Grh expression across the two genotypes. (G) Quantitative comparison of Grh levels in the larval Type I NSCs in control and heteroallelic combination (*grh^Df^/grh^DD^*). (H-H’) Shows the terminal (A8-A10) segments for late L3 (LL3) stage larval ventral nerve cord (VNC). Compared to heterozygous controls (*grh^Df/+^*)(H), which show no surviving NSCs in the late L3 stage, *grh^Df^/grh^DD^*(H’) shows a block of NSC apoptosis. (I) Shows the graphical comparison of the surviving NSCs in A3-A10 segments of late L3 stage VNC. (J-L) MARCM clones for NSCs with *grh^DD^* (J) show a reduction in Grh levels (K) and decreased cell proliferation (L) in clones of thoracic NSCs in late L3 (LL3) stage. Cyan arrowheads indicate a clone of mutant NSCs marked by GFP, which show a reduction in Grh expression; pink arrowheads point to a control NSCs (not marked by GFP), showing no change in Grh expression. NSCs (NBs) are marked with Deadpan (Dpn). Scale bars for C-D, G-G’ are 30 µm, and E-F, H and J are 10µm, respectively.

In case of *grh^DD^/grh^Df^* the expression of Grh was also reduced from Type II NSCs and their associated INPs as well (Fig S6G and S6H), and the mild developmental delay observed in this case was reminiscent of the delay seen in the case of trans-heterozygotes for *grh* deficiency and *grh^370^*(a hypomorphic allele of CNS specific O-isoform of *grh*) (*grh^370^/grh^Df^)* [26]. Therefore, we also checked the trans-heterozygotes of *grh^DD^* and *grh^370^* allele (*grh^DD^/grh^370^)* and observed a reduction in the levels of Grh from NSCs of larval CNS (Fig S6A-S6D) without any change in the levels of Grh in the wing disc (Fig S6E). This result further corroborated the NSC-specific activity and function of the deleted enhancers.

Next, we checked if the reduction in the levels of Grh in CNS owing to the double deletion had any discernible impact on cell physiology. Here we observed a proliferation defect (Fig 3J-3L) and a partial block of Hox-dependent NSC apoptosis in abdominal and terminal (A3-A10) segments of the CNS (Fig 3H and 3I), which are characteristic mutant phenotypes reported for *grh* gene in developing CNS [26].

Collectively, our result underlined that the robustness of *grh* expression relies on multiple enhancers, and different combinations of these enhancers may regulate its expression in various subsets of cells. The deletion experiments further established that the enhancer redundancy results in phenotypic robustness of *grh* expression which is a consequence of the cumulative activity of multiple enhancers. Lastly, these results emphasised that an enhancer may show activity in a specific tissue; however, the relevance of this activity (in cell physiology and tissue-specific gene expression) may need functional testing by deleting the enhancer in a sensitised background.

### Epigenetic status of *grh* and its CREs in larval NSCs, GMCs and neurons

Grh protein is known to be expressed in larval NSCs and GMCs but not in the neurons; this is reiterated by the RNA-Seq results for NSCs and neurons [41]. However, the epigenetic status of *grh* across these three cell types had not been elucidated so far. A previous study used a combination of Targeted DamID and NGS (TaDa-seq) to investigate the genome-wide binding profile of five representative chromatin-binding proteins across the three cell types in larval and adult CNS. These chromatin binding proteins were **Brm** (Brahma: SWI/SNF ATPase chromatin remodeler associated with H3K27ac), **HP1a** (Heterochromatin Protein 1a: a transcriptional repressor which is a reader for H3K9me3), **Pc** (Polycomb: a transcriptional repressor which is a reader for H3K27me3, representing PcG chromatin), Histone **H1** (a linker histone, found in repressive chromatin) and the core subunit of **RNA Pol II** (RNA Polymerase II: indicating permissive and actively transcribed chromatin) [7].

An underlying motivation to identify and functionally validate NSC-specific enhancers (*Int-2BF1* and *Int-2C*) of *grh* was to check how the epigenetic status of a validated tissue-specific enhancer compared with that of the (*grh*) gene in a given cell type and to see how the status change across the three cell types (NSCs, GMCs and neurons). To this end, we analysed previously published TaDa-seq data (Targeted DamID-seq) [7], and observed that the *grh* and the two deleted CREs show marks for RNA Pol II and Brm in the larval NSCs (Fig 4A and 4A’). In the case of GMCs, while the Brm marks were present on *grh* and CREs, RNA Pol II showed very little occupancy on the two (Fig 4B and 4B’ vs 4A and 4A’). In the case of the neurons, the *grh* and the two CREs did not show any occupancy by Brm and RNA Pol II (Fig 4C and 4C’). For the repressive marks, we observed that NSCs and GMCs show Pc and HP1a marks, and H1 showed very low occupancy on the *grh* locus and the two CREs (Fig 4A-4A’ and Fig4B-4B’). This suggests that *grh* was active in some of the NSCs and was repressed in others. This, however, changed in the case of neurons, where significant H1 and HP1a repressive marks were observed on *grh* and the two CREs (Fig 4C and 4C’), which is consistent with the fact that *grh* is not expressed in neurons in postembryonic stages.

**Fig 4.**
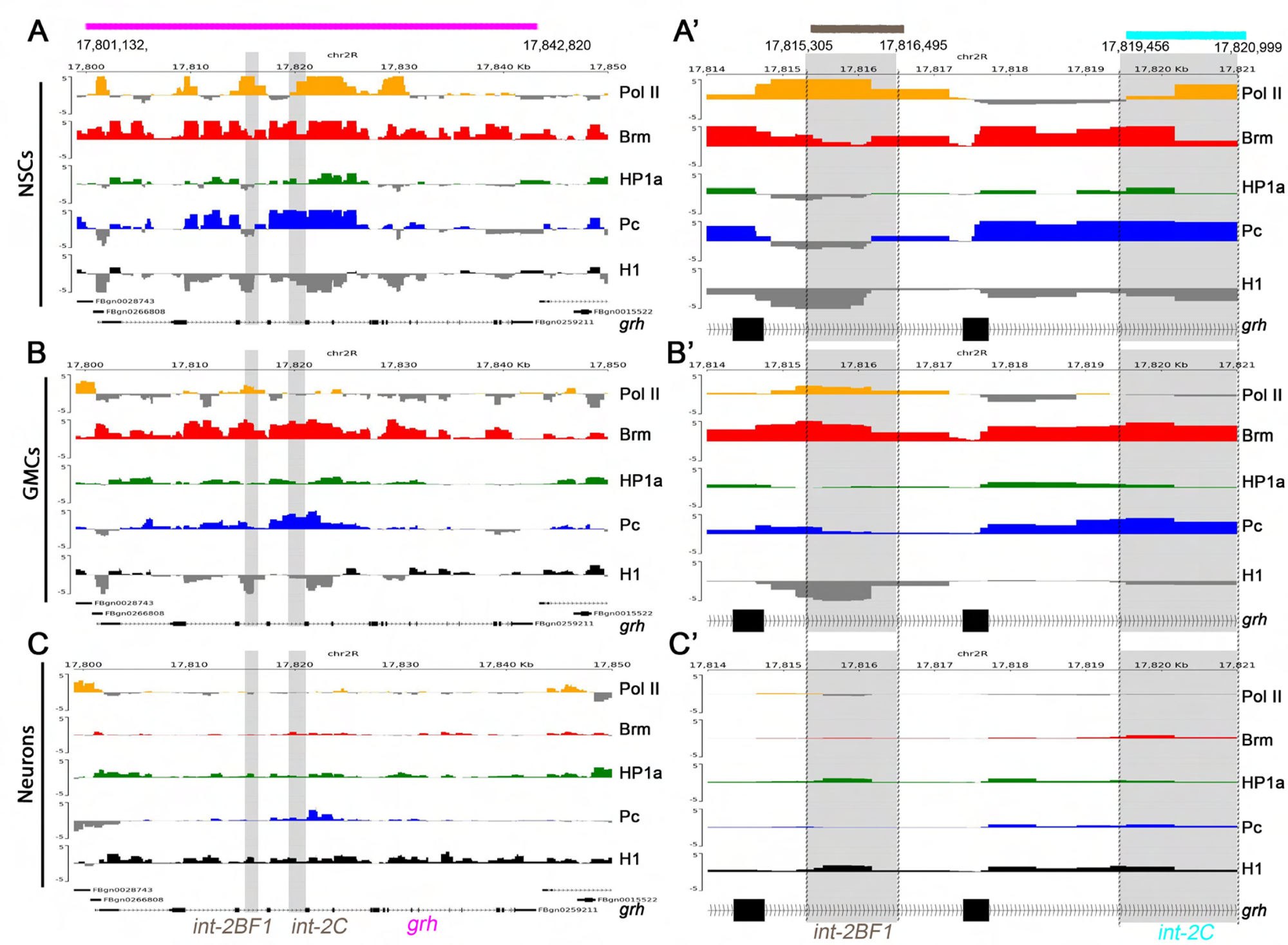
Epigenetic status of *grh* and its tissue-specific CREs are in sync in the NSCs, GMC and neurons. (A-C) Show occupancy profile of five chromatin marker proteins on *grh* gene and its enhancers *Int-2BF1* and *Int-2C* (A’-C’) in NSCs, GMCs and neurons. The five chromatin markers are RNA Pol II (orange), Brm (red), HP1a (green), Pc (blue) and H1 (grey). The binding profiles suggest that the epigenetic status of CREs is in sync with that of the whole *grh* gene in the NSCs, GMC and neurons. The position of *Int2BF1* and *Int-2C* are indicated in the context of the whole *grh* gene in A, B and C panels by grey shading. Y-axis limits are set from −5 to 5 for occupancy comparison across the tracks. Coloured horizontel lines above the panels show the genomic extent of *grh* gene (from Chr-2R: 17,801,132 to 17,842,820, pink), *Int-2BF1* (Chr-2R: 17,815,305 to 17,816,495, brown) and *Int-2C* (Chr-2R: 17,819,456 to 17,820,999, cyan). The coordinates indicated at the top are in kilobases.

Importantly, RNA Pol II binding on *grh* indicated that it was actively transcribed in NSCs but not in GMCs and neurons, suggesting that *grh* transcripts or protein was most likely inherited by GMC from the NSCs. To experimentally test this, we did a smFISH (single molecule Fluorescence Insitu Hybridisation) for *grh* transcripts in the cells of larval CNS using *grh*-specific probes. We observed that *grh* mRNA was confined primarily to NSCs and was not seen in GMC or neurons (Fig 5A). To further test this, we analysed the expression of Grh in the GMCs associated with the NSCs, which show a complete loss of Grh expression in *grh^DD^/grh^Df^* (pink arrowheads, Fig 5C’’). Here, we observed that these GMCs did not express Grh like the NSCs (yellow arrowheads in Fig 5B’’ vs 5C’’). This would not be the case if GMCs were independently transcribing *grh,* reiterating that the GMCs inherit the Grh protein from the NSCs.

**Fig 5.**
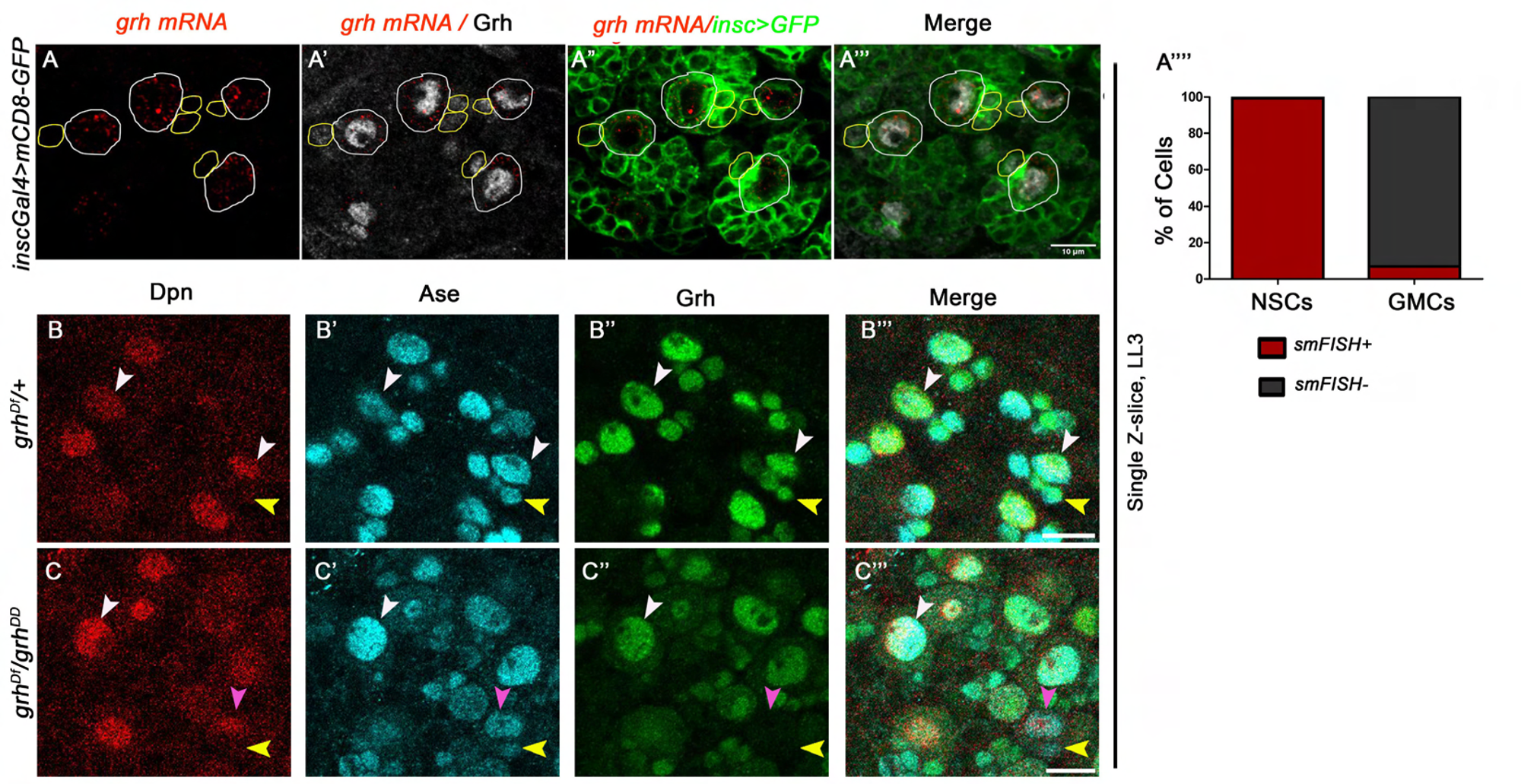
*grh* is transcribed in NSCs but not in GMCs. (A-A’’’’) smFISH detects *grh* mRNA in NSCs but not in GMC associated with it, or in other surrounding cells. Grh antibody is used to mark NSCs and GMCs. (A’’’’) Graphs showing the % of cells positive for smFISH signal (for *grh* transcripts) in NSCs and GMCs. NSCs (NBs) and GMC are marked by white and yellow circumferences, respectively. (B) Comparison of the Grh expression in GMCs associated with the thoracic NSCs for heterozygous (*grh^Df^/+*: control) larvae versus heteroallelic (*grh^Df^/grh^DD^*) larvae (C). The subset of NSCs (Dpn^+^, Ase^+^) that showed a complete loss of Grh expression in the heteroallelic (*grh^Df^/grh^DD^*) larvae also showed a complete loss of Grh from the GMCs (Dpn^-^, Ase^+^) as well, suggesting that *grh* is expressed only in NSCs and Grh protein is most likely inherited from NSCs to GMCs. White and pink arrowheads indicates NSCs, and yellow arrowheads indicates GMCs showing a complete loss of Grh expression. Scale bar is 10 µm.

Collectively, we observed that the *grh* gene was expressed only in the NSCs, and it was the Grh protein that gets inherited by GMCs, which could be a general strategy for regulating the expression of cell fate determinant proteins in intermediate progenitor cells. These results also showed for the first time in vivo that the epigenetic state of a gene and its functionally validated (deleted) tissue-specific CREs are broadly in sync with each other, and this synchrony is maintained across different cell types of the developing CNS.

## Discussion

Enhancers are the epicenters of tissue-specific gene regulation. Our study used the CNS-specific expression of the *Drosophila grh* gene to make a case for deleting the enhancers in a sensitized background of other enhancer deletion to functionally validate their role in tissue-specific gene regulation. By identifying novel enhancers and subsequently deleting two of them, we established their collective importance in regulating *grh* expression and showed that *grh* relies on multiple enhancers for its robust expression in NSCs of CNS, with different combinations of CREs playing a critical role in regulating *grh* in various subset cells. We also found that the epigenetic status of these enhancers is in sync with that of *grh* gene across the three cell types of developing CNS, and *grh* is transcriptionally expressed only in NSCs and not in GMC, which inherits the Grh protein from the former.

In our experiments, we found that single deletions of two NSC-specific CREs did not impact the Grh expression in the CNS, and both the CREs had to be deleted together to show a significant reduction in Grh expression and associated physiological phenotype. This suggested that multiple enhancers may show activity in a specific cell type, but their role in gene regulation may be context-specific and become apparent only in a sensitized background of deletion of another enhancer. Our results are in line with an extensive enhancer deletion study carried out earlier in mouse where single and double deletions were made for ten enhancers responsible for limb development. In this study, it was observed that none of the ten single enhancer deletions showed any discernible phenotype in limb development, but double deletion of enhancers resulted in limb abnormalities. This emphasised the importance of the sensitised background and cumulative activities of the functionally redundant enhancers in regulating the tissue-specific gene expression [40].

Considering the diversity of Grh expression in neural and non-neural tissues [13], we expected its regulation to be more complex and dependent on multiple CREs. This was supported by the fact that double deletion of the enhancers reduced Grh expression significantly in the majority of the NSCs and could abolish it completely from a small minority of cells. Therefore, we were surprised that only one CRE (*Ex1up*) showed activity in epithelial wing discs, and we expect that there might be other CREs regulating *grh* expression which may lie outside the genomic region bracketed by the Type I insulator elements [38]. A complex transcriptional regulation for *grh* is also underlined by a previous study in the vertebrates wherein one of the *grhl2* enhancers important for craniofacial development was deleted; however, no physiological phenotype was observed despite a 50% decrease in RNA transcripts [34]. This emphasised the robust and multi-modular transcriptional networks regulate Grhl2 expression in the vertebrate system, similar to what we find in the case of *Drosophila grh* as well.

Overall reduction in the levels of Grh in the CNS that we observed in the case of *grh^DD^/grh^Df^*can be interpreted in the light of the Source-Sink model used to explain epigenetic regulations of gene transcription [42]. In this model, genomic DNA operates as the “Sink”, and nucleosomes and epigenetic factors function as the “Source.” The model suggests that different source proteins compete for binding on the sink/DNA, thereby affecting genomic equilibrium and gene regulation in a cell. An alternative way to think about it is the converse, where changing the sink availability could alter the overall free source levels. Now in NSCs expressing Grh the free levels of repressive factor in nucleus would normally be negligible, and the activating factors bind and maintain the Grh protein expression at a certain level. In this background, the deletion of one enhancer does not affect this balance sufficiently, but the deletion of two enhancers increases the free repressor levels beyond a threshold, thereby interfering with the binding of the activating factors on the DNA and thus reducing the *grh* expression in majority of the NSCs. The few cells where Grh expression is completely abrogated are the cells where these two deleted enhancers are critical for maintaining active gene expression.

The genome-wide binding profile of the five representative chromatin-binding proteins showed Pc and HP1a marks on *grh* in NSCs, which suggest that all the NSC lineages may not express Grh simultaneously, which corroborated with our previous finding that the NSCs in the terminal region (in A8-A10 segments) show a temporal delay in initiating Grh expression compared to their thoracic and abdominal counterparts [31]. This analysis also showed that *grh* is transcribed only in NSCs and not in GMC, and Grh protein is inherited from NSCs to GMCs, which could be a general strategy to regulate the expression of cell fate determinant proteins in intermediate progenitor cells. Escargot protein, which is expressed in *Drosophila* intestinal stem cells and intermediate progenitor cells (like enteroblast and enteroendocrine cells), but is repressed in the differentiated progeny [43–45] may be employing a similar mechanism.

Our analysis also shows that tissue-specific CREs are broadly in sync with the epigenetic state of the *grh* gene in all three cell types of the CNS. Such epigenetic synchrony is often proposed for tissue-specific gene regulation in vivo but has not been shown experimentally in a developing organism. More importantly, the enhancers being analysed were not functionally validated by their deletion. In this work, we first established the importance of the enhancers by deleting them and thereafter showing that the epigenetic status of these enhancers is in synchrony with that of the *grh* gene across different cell types. However, it remains to be tested experimentally how the epigenetic status of the gene changes when these tissue-specific enhancers are deleted. The double deletion generated by us for *grh* will be instrumental in addressing this question in future. Lastly, considering the role of Grh as a tTF, it will be interesting to investigate the effect of the double deletion of CREs on the generation of cellular diversity in CNS.

## Materials and methods

### Fly stocks and fly husbandry

The following fly lines were used in this study: *Canton-S* (BDSC-64349), *UAS-dcr2; inscGAL4-UASmCD8-GFP* [46], *Notch^RNAi^* (BDSC-28981), *PntGal4-UASmCD8-GFP* (Jurgen Knoblich, IMP, Vienna, Austria), *grh GAL4* (BDSC-64345). Egg collections were done for 4 hr, and flies were grown at 25°C until the desired time points of dissection. The L1 and L2 stages were divided into three 8 hour intervals to define the early, mid, and late stages; for L3, this interval was 16 hours. The age was calculated as the hours after egg laying (AEL).

### Generation of transgenic flies

*Ex1up-lacZ*, *Int-1-lacZ*, *Int-2A-lacZ, Int-2BF1AB-3X-lacZ*, *Int-2C*-*lacZ*, *Int-3-lacZ*, *Int-7-lacZ*, *Int-8-lacZ, Int-9-lacZ*, *Int-11-lacZ, Int-14-lacZ* transgenic lines are generated using hs43-nuc-lacZ vector [47] modified for by site-specific insertion (referred to as *reporter-lacZ* here). All the constructs were inserted at attP2-68A4 on Chr-3L using phiC31-based integration system [48].

*Int-2BF1A-lacZ*, *Int-2BF1B-lacZ*, *Int-2BF1A*’*-lacZ*, *Int-2BF1*B’*-lacZ*, *Int-2BF1AB-lacZ*, *Int-2BF2-lacZ*, *Int-2BF3-lacZ* were generated using hs43-nuc-lacZ vector [47] by P-element-mediated transformation.

The enhancers were PCR amplified using region-specific primers and cloned into appropriate *reporter-lacZ* vectors.

### Immunohistochemistry and image acquisition

Brains of the desired larval stages were dissected and fixed in 4% paraformaldehyde in 1x PBS containing 0.3% TritonX-100 for 30 min at room temperature (RT) and followed by washing with 1X PBS containing 0.3% Triton X-100 for 30 minutes at RT. Then the brains were immunostained with primary antibodies overnight at 4°C. Embryos of stage 16^th^ were stained as previously described [49]. The following primary antibodies were used: rabbit anti-Dpn (1:5,000; Bioklone, Chennai), rabbit anti-Grh (1: 2,000; Bioklone), chicken anti-β-gal (1: 2,000, ab9361, Abcam), Ase (CDFD animal house, 1:1000), Grh Exon 4+5(this study, CDFD animal house, 1:1000). Secondary antibodies conjugated to Alexa fluorophores from Molecular Probes were used: Alexa Fluor 405 (1:250), Alexa Fluor 488 (1:500), Alexa Fluor 555 (1:1,000), and Alexa Fluor 647 (1:500). The samples were mounted in 70% glycerol. Images were acquired with Zeiss LSM 900 confocal microscope and processed using ImageJ and Adobe Photoshop CS3. Scale bars are in µm and mentioned in the figure legends. White and pink arrowheads point NBs in all the figures; GMCs are indicated by yellow arrowheads. All the experiments were repeated at least 3 times, with controls and tests processed and imaged simultaneously in the same settings. All the images are representative of a minimum of 7 CNS specimens analysed.

### smFISH-IF protocol

The smFISH was performed by following the protocol described in literature [50]. BioSearch probe designer was used to design the probes for *grh* gene against the coding sequence. 20nt probes were generated and checked for the off-targets. Generated probe (total 47) sequences were sent to Biosearch Technologies for synthesis with Fluorophore Quasar 570. In brief, larval CNS were dissected from 3^rd^ instar larvae in 1xPBS and fixed in 4% paraformaldehyde in 1x PBS containing 0.3% TritonX-100(PBSTx) for 30 min at room temperature (RT) and followed by 3 quick washes with 1X PBS containing 0.3% Tween-20 (PBST). Brains were washed 3 times for 15 minutes each at 25^ο^C with PBST. Then, brains were incubated in wash buffer (10%v/v deionised formamide 1 in 2x saline sodium citrate (SSC)) solution for 5 mins at 37 ^ο^C. Then, brains were incubated in hybridization buffer (10% v/v deionized formamide, 10% v/v of 50% dextran sulphate solution (final dextran concentration = 5%) in 2x saline sodium citrate (SSC) solution) with the smFish probe concentration 0.5µm at 37^ο^ C for 8–15 hours with gentle rocking. For protein immunofluorescence, mouse anti-Grh (1:2000, Bioklone) was added in this step and incubated overnight. Thereafter, brains were rinsed 3 times in wash buffer. After this, brains were washed 3 times for 15 minutes each with wash buffer at 25^ο^ C. Next, brains were incubated with secondary antibody against Grh (Alexa Fluor 647) for 1 hour and 30 minutes at 25° C. Finally, brains were washed 3 times with wash buffer for 10 minutes each. The samples were mounted in 70% glycerol. Images were acquired with a Zeiss LSM 900 confocal microscope. Twelve specimens were analysed across three repetitions of the experiment, and a representative image is shown.

### CRISPR-Cas9 deletion of Grh enhancers

The individual deletions of the *Int-2BF1* and *Int-2C* were performed using a double gRNA/Cas9-based strategy as described in [4]. The following guide RNA sequences were chosen:

Left guide for *Int-2BF1* deletion: GCATATGCCCCGCAGATCC

Right guide for *Int-2BF1* deletion: GGTAGTACTGTGAGAATTAC

Left guide for *Int-2C* deletion: GAGGAATATATGACATTAGT

Right guide for *Int-2C* deletion: GTGCAAAGGAACCCATCATGT

Both the left and Right guides for both the enhancers were cloned into a double gRNA vector where each of the guides is expressed under its own U6 promoter [4]. The constructs were then integrated into the attP40 landing site on the second chromosome. The two gRNA-containing males were then crossed to females of nos-cas9, and the resultant offspring containing both the gRNA and the cas9 were obtained. These were then crossed to second chromosome balancer flies, and 25 independent stocks were subsequently established. These stocks are homozygosed and flies from each stock were tested for the presence or absence of deletion by PCR. Two independent stocks containing the desired deletion were obtained, and PCR mapped the breakpoints. The following primer pairs were used to screen for the deletion:

Primers for *ΔInt-2BF1* screening

*Amplicon A*: *CTGTACTCTGCAAGTGGTCC* and *TCGCTCTAGGTACATACCCTG*

*Amplicon B*: *AACCATCACACTTTGTCTGTC* and *GTATTTGTACGAGCATATGC*

Primers for *ΔInt-2C* screening

*Amplicon A*: *CAATTGGGCACATCGAATGGA* and *TCTGGCCATTGAGCTTTCC*

*Amplicon B*: *CAGTAACGCTCGGAATTTGC* and *ATCCTTGCGCAAGTCTTCG*

For the generation of the double deletion of CREs (*ΔInt-2BF1, ΔInt-2C*), females from fly strain Δ*Int-2BF1/SM6 Cyo; nos-cas9/TM6B,Tb* was crossed with the *Int-2C* double gRNA containing males and similar steps were followed as mentioned for the generation of single deletions. 70 individual lines were established and screened for deletion by PCR, and only 1 line tested positive for deletion of both the enhancers.

Primers for double mutant (*ΔInt-2BF1, ΔInt-2C*) screening

*Int-2BF1 Amplicon*:

*AACCATCACACTTTGTCTGTC* and *GTATTTGTACGAGCATATGC*

*Int-2C Amplicon*:

*CAATTGGGCACATCGAATGGA* and *TCTGGCCATTGAGCTTTCC*

Activity interaction experiments shown in Fig 2 were done with the following genotypes:

*w*; *grh*Δ*Int-2BF1/grh*Δ*Int-2BF1; Ex1up-lacZ-68A4/+*

*w*; *grh*Δ*Int-2BF1/grh*Δ*Int-2BF1; Int-2C-lacZ-68A4/+*

*w*; *grh*Δ*Int-2C/grh*Δ*Int-2C; Ex1up-lacZ-68A4/+*

*w*; *grh*Δ*Int-2C/grh*Δ*Int-2C; Int-2BF1-lacZ-68A4/+*

### DamID-seq data processing

To study the chromatin state of *grh* in NSCs (neuroblasts-NBs), GMCs and neurons, bedgraphs were taken from previously published data [7] of different chromatin states (GEO GSE77860). Scaled averaged bedGraphs were downloaded as each dataset is divided by its standard deviation suitable for better comparison as described in the study.

All the five chromatin binding proteins were visualised in Integrated Genomic Viewer IGV for the binding on *grh* and its enhancers in the three datasets (NB or NSCs, GMCs, neurons). The DamID tracks are generated using vector software pyGenomeTracks [51, 52].

## Data Availability Statement

All relevant data are within the manuscript and its Supporting Information files.

## Supplementary Figures

**Fig S1.**
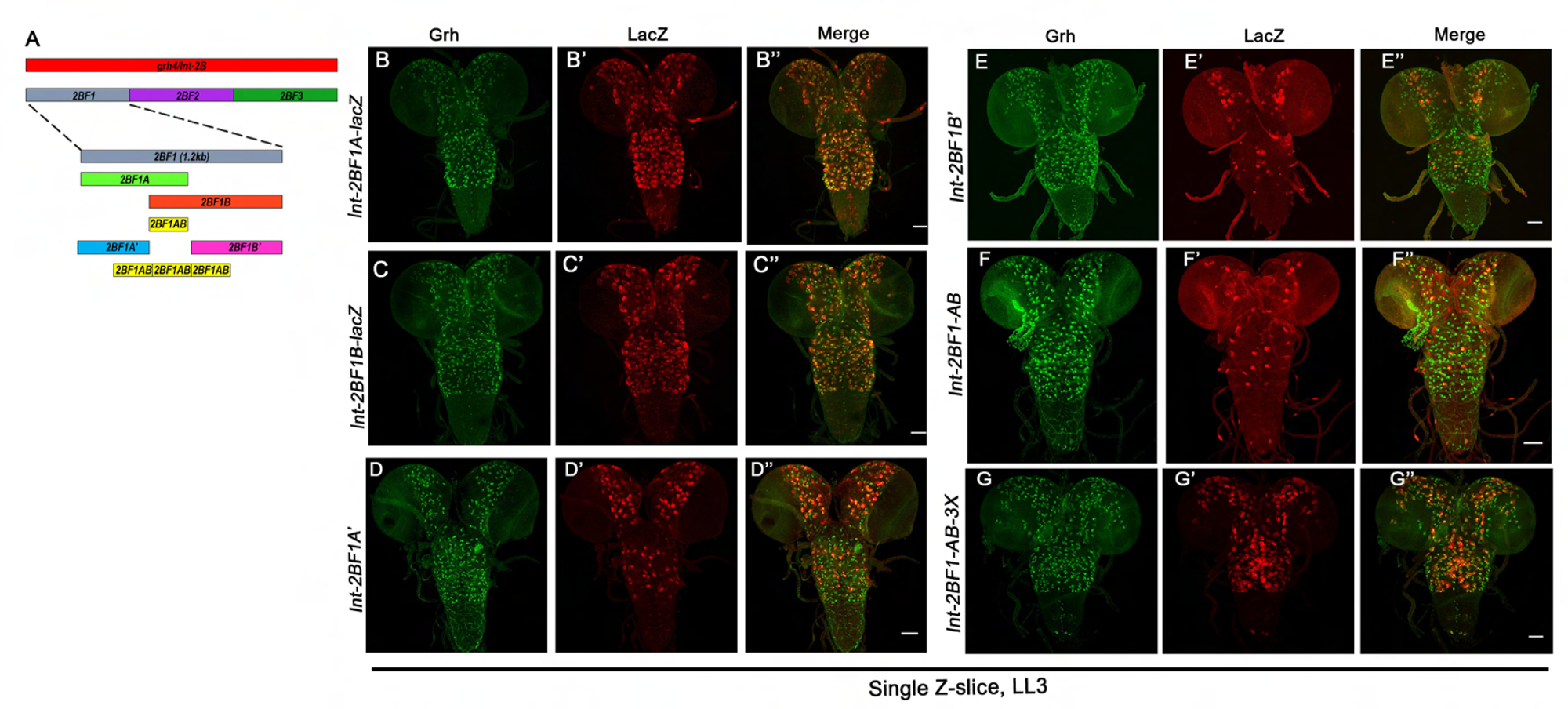
Sub-fragmentation of 4Kb *grh enhancer*. (A) Schematic for fragmentation of 4 Kb *grh* enhancer (*grh4/Int-2B*). (B-C) Subfragments of *Int-2BF1* (*Int-2BF1A* and *Int-2BF1B)* show activity in most Grh expressing NSCs. (D) *Int-2BF1A’-lacZ* is expressed strongly in a subset of Grh-expressing NSC of the central brain, and its expression in the VNC is highly reduced. (E) *Int-2BF1B’-lacZ* could be visualised only in a subset of Grh expressing NSCs of the central brain and thoracic region of VNC. (F) The 150 bp overlap between *Int-2BF1A* and *Int-2BF1B* is expressed only in very few central brain cells and VNC cells. (G) The trimerised version of the 150 bp overlap *(Int-2BF1AB-3X-lacZ)* is expressed in the majority of the Grh-expressing NSCs in the late L3 stage CNS. Scale bars are 50µm.

**Fig S2.**
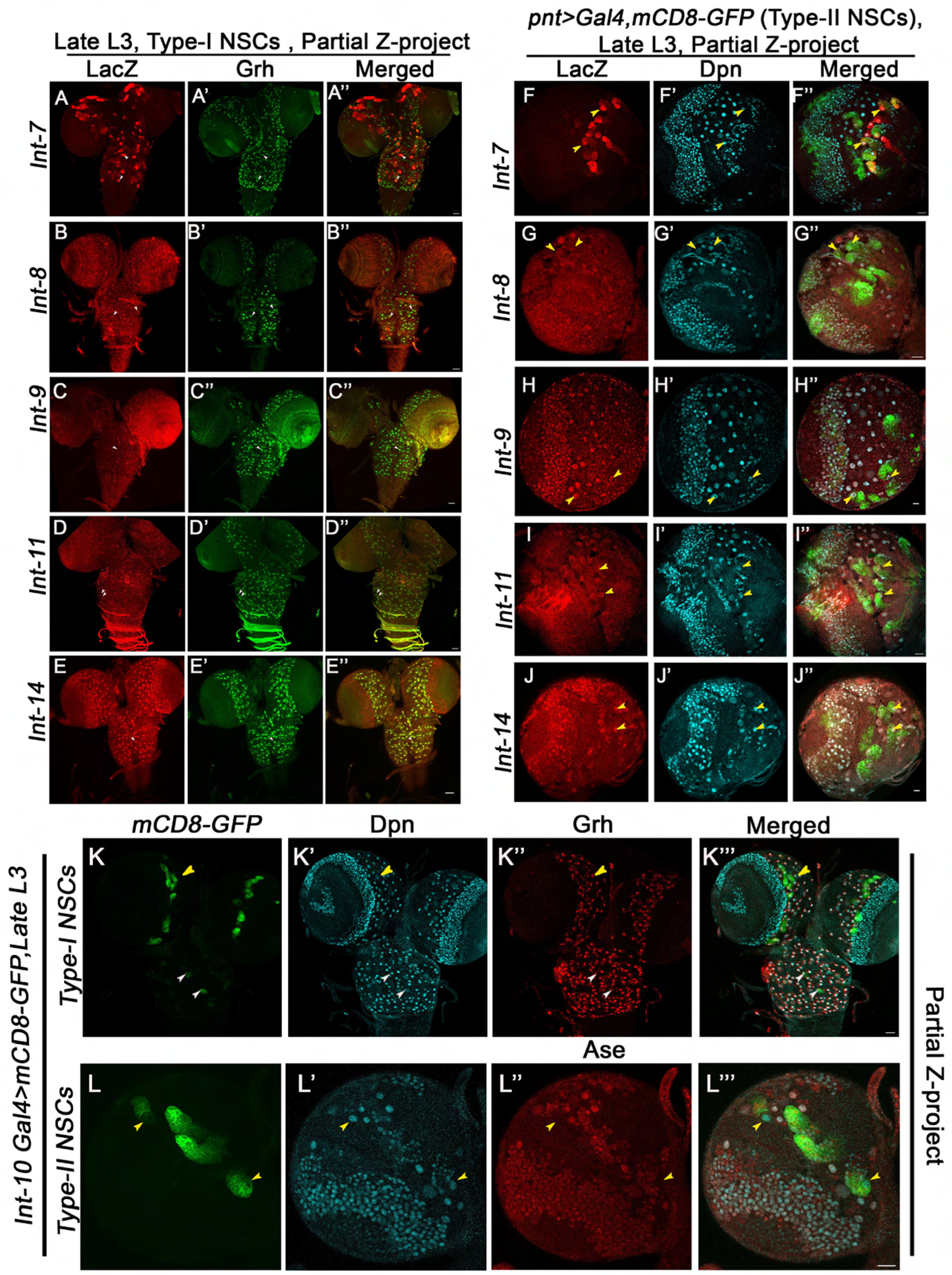
Expression of *grh* CREs in larval CNS. (A-J) Shows the varying levels of activity of various *grh enhancer-lacZ* reporters (*Int-7, 8, 9, 11*, and *14*) in Type I NSCs (A-E) and Type II NSCs (F-J) in late L3 stage of larval CNS. *Pnt-GAL4>UAS mCD8-GFP* is used to mark the Type II NSCs (in panels F-J)*. Int-7* showed some cells with very high and some with very low activity in NSCs. *Int-8, 9, and 11* showed very weak activity in Type-I NSCs and Type-II NSCs. *Int-14* showed significant expression in both Type I and Type II NSCs. (K-L) Shows the expression of *Int-10-GAL4>UAS-mCD8-GFP* (BDSC line 64345) in Type I and Type II NSCs of late L3 stage larval CNS. *Int-10-GAL4* is strongly expressed in Type-II NSCs and showed a very weak and limited expression in Type-I NSCs. NSCs are marked with either Dpn or Grh or Asense. White arrowheads indicate Type I NSCs (NBs) and yellow arrowheads indicate Type II NSCs. Type II NSCs in panels L are identified by Dpn^+^/Ase^-^ staining. Scale bars are 20µm.

**Fig S3.**
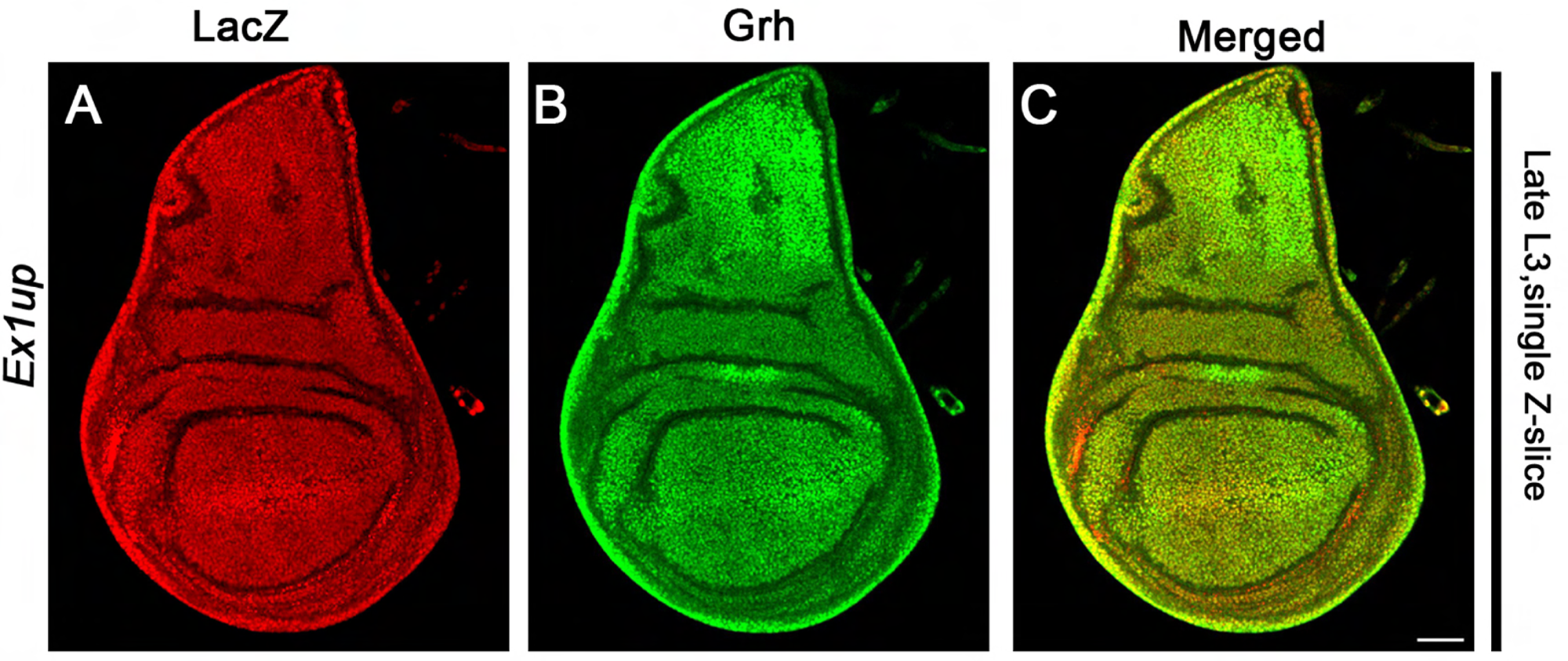
Expression of grh CREs in Wing Disc. (A-C) Shows the expression of *Ex1up-lacZ* in wing discs of the late L3 larvae. Scale bars are 20µm.

**Fig S4.**
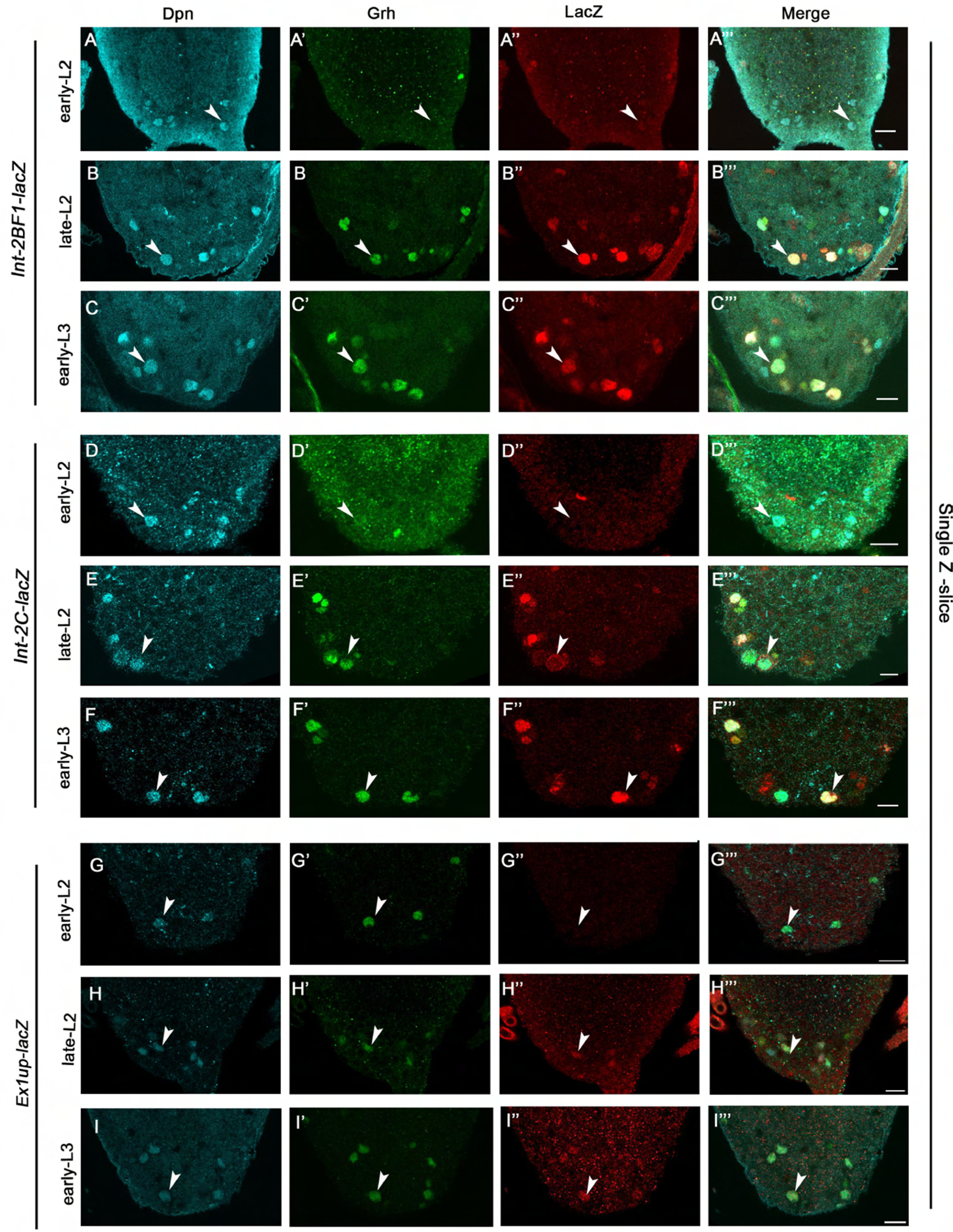
*Int-2BF1, Int-2C* and *Ex1up-lacZ* activity in A8-A10 NSCs recapitulates the temporal increase of Grh seen from early L2 to early L3 stage. (A-I) Shows *Int-2BF1, Int-2C* and *Ex1up-lacZ* activity in A8-A10 NBs in early L2, late L2, and early L3 stages of development, NSCs are marked with Dpn. White arrowheads indicate the expression of lacZ in terminal NSCs. Scale bar is 20µm.

**Fig S5.**
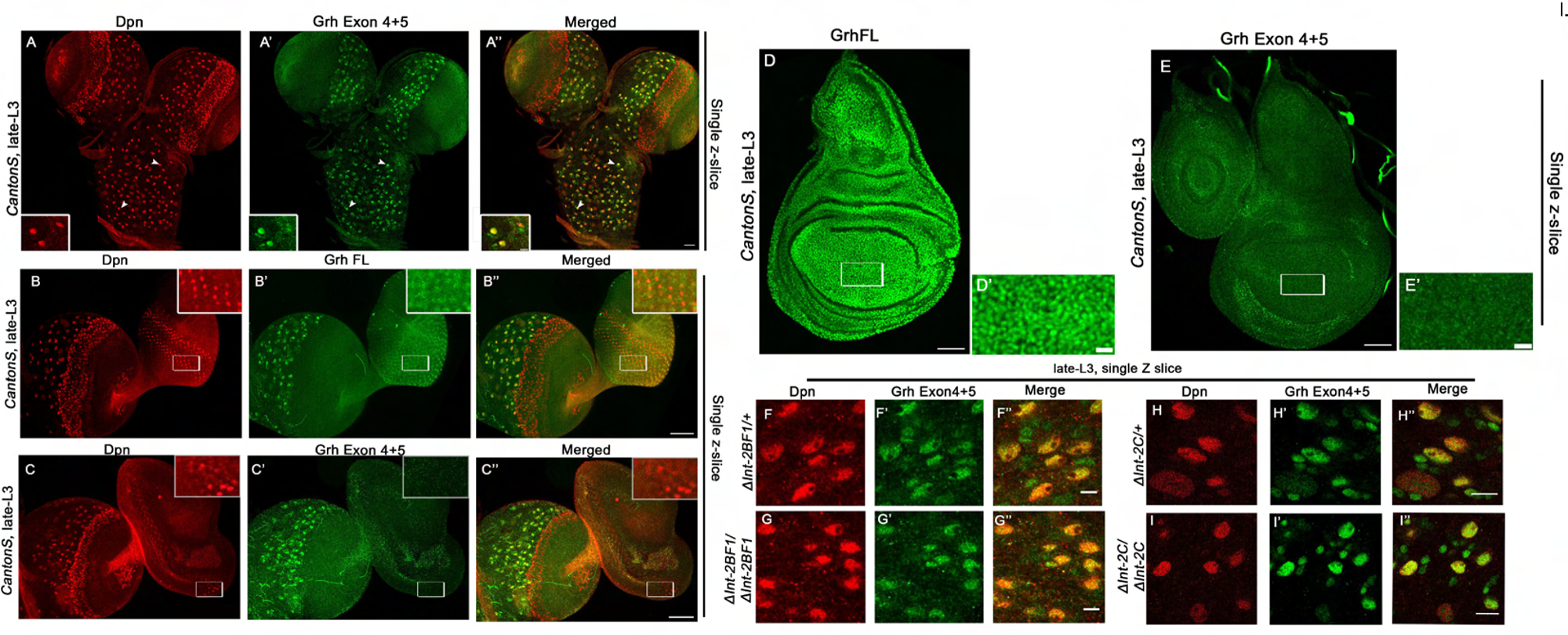
The expression of CNS-specific GrhO isoform is unaltered in the single deletions for *Int-2BF1* and *Int-2C*. (A-A”) GrhO isoform-specific antibody (Grh Exon4+5) (raised against 205 residues from Exon4+5) marks both NSCs and GMCs in late L3 larval stage CNS of Canton S larvae. (B-C) Compares the staining of the antibodies raised against full-length Grh protein (GrhFL-detecting all the isoforms of Grh) and 205 residues from Exon4+5 (specific for GrhO isoform, Grh Exon4+5) in late L3 larval central brain and eye disc of Canton S larvae. GrhFL antibody shows expression in both CNS and eye discs, while GrhO (Grh Exon4+5) specific antibody detects Grh exclusively in CNS but not in the eye discs. (D-E) Compares the staining of Grh FL and GrhO specific (Grh Exon4+5) antibody in late L3 stage larval wing disc of Canton S larvae. (F-I) Shows that the expression of GrhO is unaltered in the heterozygotes and homozygotes for the single deletion of *Int-2BF1* (F-G) and *Int-2C* (H-I) in the thoracic NSCs of late L3 stages CNS. NSCs (NBs) are marked by Deadpan (Dpn). Scale bars are 20 µm (A-C), 50 µm (D-E), 20 µm (D’-E’), 10 µm (F-I).

**Fig S6.**
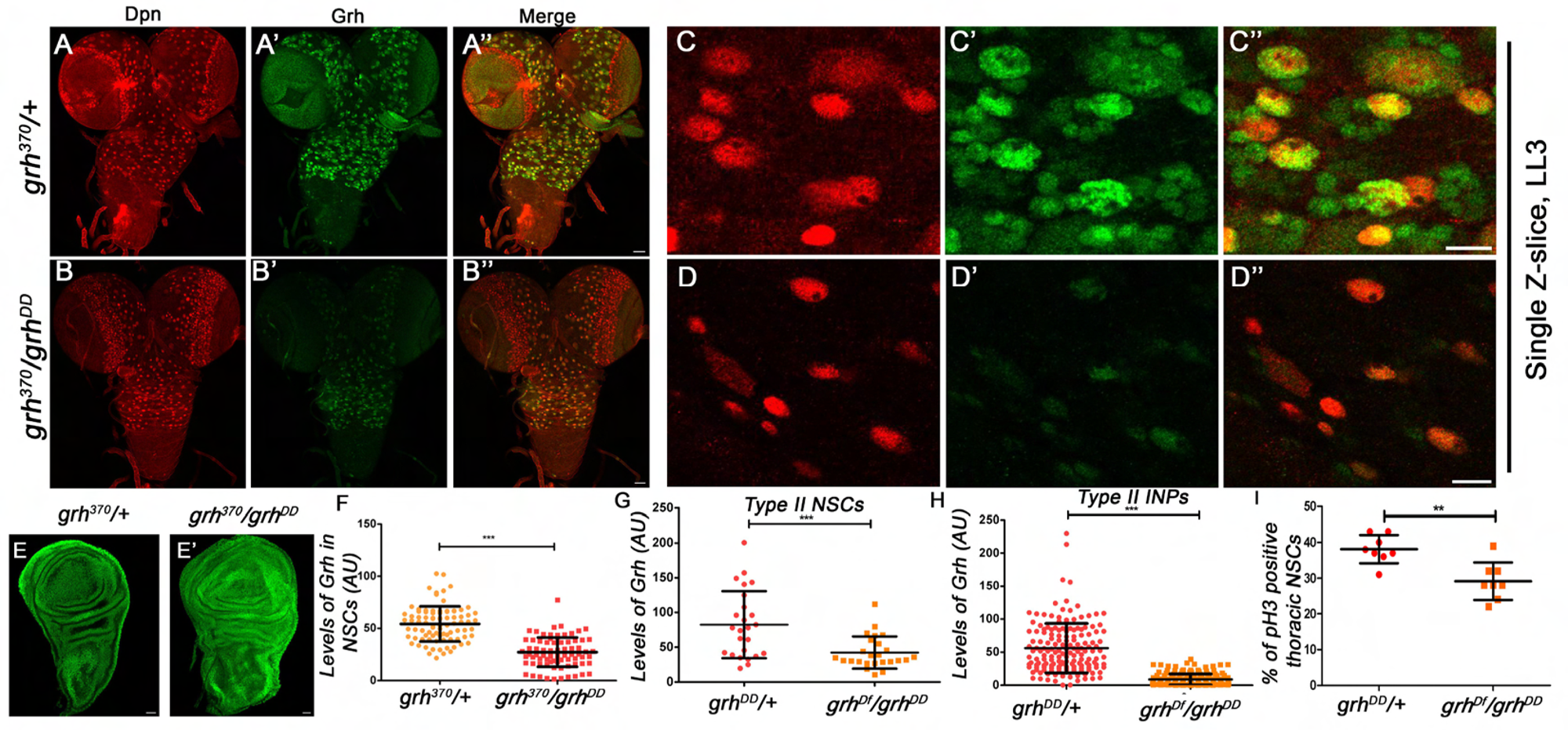
Trans-heterozygotes for the GrhO specific allele (*grh^370^*) and Double deletion of *Int-2BF1* and *Int-2C* enhancers (*grh^DD^*) compromise Grh expression, specifically in NSCs. (A-D) Compares the Grh staining in the late L3 stage CNS and thoracic NSCs of heterozygous *(grh^370^/+)* and heteroallelic combination (*grh^370^/grh^DD^*). The heteroallelic combination shows a dramatic decrease in Grh expression in the NSCs, with some NSCs showing a complete loss of Grh (similar to what is seen for *grh^Df^/grh^DD^*), while the wing discs (E-E’) do not show any change in Grh expression across the two genotypes. (F) Quantitation of Grh intensity in the larval NSCs of heterozygous *(grh^370^/+)* and heteroallelic combination (*grh^370^/grh^DD^*). (G-I) Shows comparison of levels of Grh in Type-II NSCs (G), associated INPs (H) and mitotic index in thoracic NBs of heterozygous (*grh^DD/+^)* vs heteroallelic (*grh^Df/DD^*) combinations of *grh.* NSCs (NBs) are marked with Deadpan (Dpn). Scale bars for A-B, E and E’ are 30 µm, and C-D are 10 µm.

**Fig S7.**
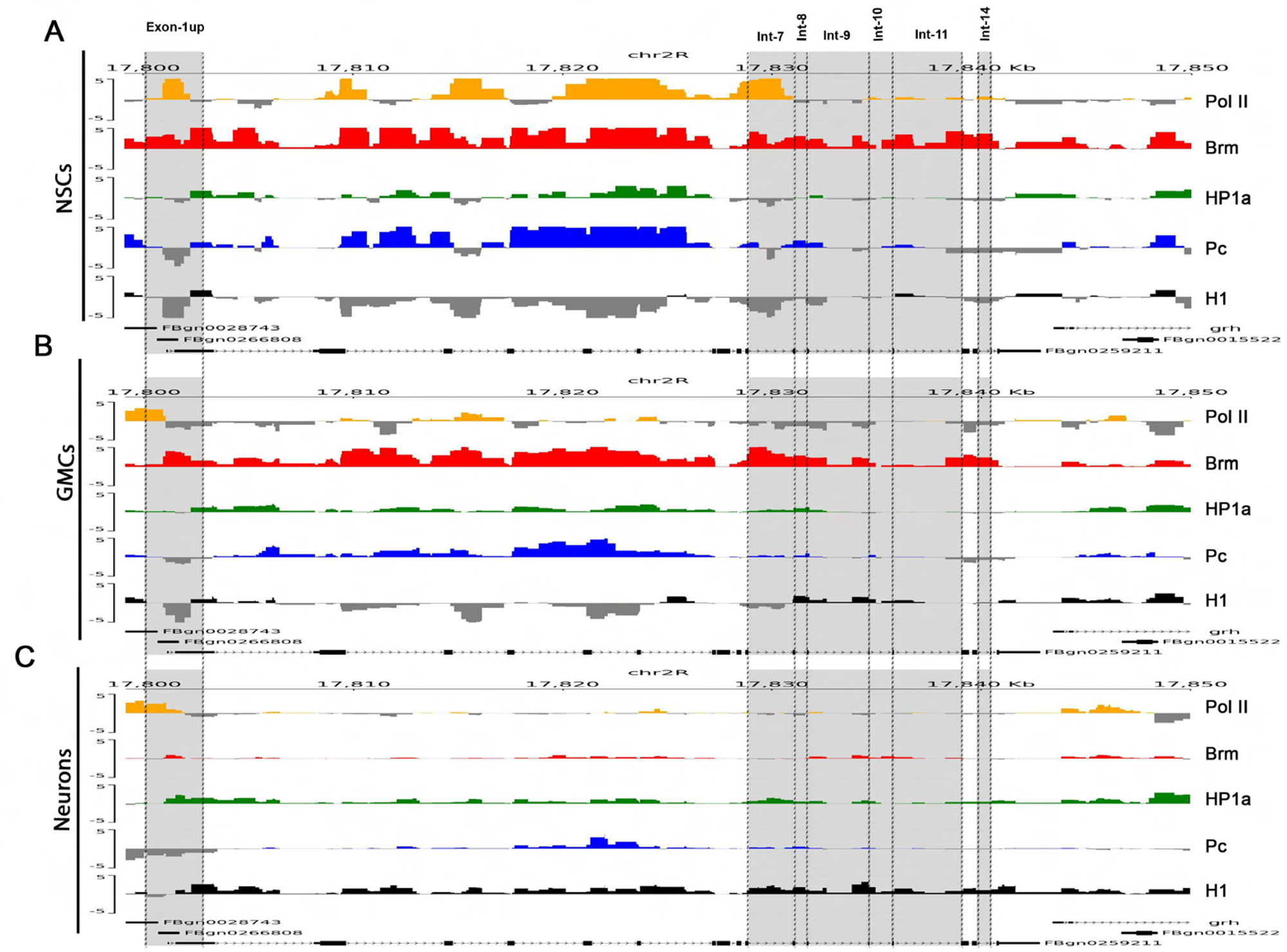
Pygenome tracks for NSC, GMC and neurons showing the relative positions and chromatin marks for all the enhancers analysed for *grh* gene. The details of the epigenetic marks for each the enhancers are given in Table III.

## Acknowledgements

We thank R. Mann, J. Knoblich, S. Small, R. Rikhy, A. Ratnaparkhi, G. Ratnaparkhi, for various reagents and/or advice with experiments; D. Trivedi for her constant guidance and advice with the gRNA designs; BDSC, NIG-Fly, and DGRC -Japan stock centres for fly lines; DGRC -Japan for dual gRNA CRISPR-Cas9 constructs, CDFD animal facility, Bioklone Biotech Pvt. Ltd., Chennai and the Developmental Studies Hybridoma Bank at The University of Iowa for antibodies, Sophisticated Equipment Facility at CDFD for DNA sequencing and TFF at NCBS-CCAMP, Bangalore for transgenic flies. We thank Ch. Gangi Reddy for his advice and help with NGS data analysis, and C. S. Singh for his assistance in various phases of the project. This study was funded by; Department of Science and Technology, India (CRG/2021/003275); Department of Biotechnology, India (BT/PR41306/MED/122/ 259/2020; BT/PR45460/MED/12/952/2022); Wellcome Trust DBT India Alliance, India (Ref: 500171/Z/09/Z), CDFD core funds; and ICMR, India (award to R.S.) [ICMR Ref.No:3/1/3/JRF-2012/HRD-63 (40260)], CSIR, India (award to YR) [File No:09/724(0132)/2018], UGC-JRF (award to JB) [UGC-No: F.16-6(DEC.2016)/2017(NET)].

## Author contribution

RJ conceptualised the study; RS, YR, JB, and PA did the experiments; RJ, RS, YR, and JB analysed the data; PA did the NGS data analysis; VK generated the double deletion. RJ wrote the manuscript.

## References

1. Horvath, P. and R. Barrangou, CRISPR/Cas, the immune system of bacteria and archaea. Science, 2010. 327(5962): p. 167–70.

2. Mali, P., et al., RNA-guided human genome engineering via Cas9. Science, 2013. 339(6121): p. 823–6.

3. Jinek, M., et al., RNA-programmed genome editing in human cells. Elife, 2013. 2: p. e00471.

4. Kondo, S. and R. Ueda, Highly improved gene targeting by germline-specific Cas9 expression in Drosophila. Genetics, 2013. 195(3): p. 715–21.

5. Gratz, S.J., et al., CRISPR-Cas9 Genome Editing in Drosophila. Curr Protoc Mol Biol, 2015. 111: p. 31 2 1–31 2 20.

6. Southall, T.D., et al., Cell-type-specific profiling of gene expression and chromatin binding without cell isolation: assaying RNA Pol II occupancy in neural stem cells. Dev Cell, 2013. 26(1): p. 101–12.

7. Marshall, O.J. and A.H. Brand, Chromatin state changes during neural development revealed by in vivo cell-type specific profiling. Nat Commun, 2017. 8(1): p. 2271.

8. Bray, S.J., et al., Embryonic expression pattern of a family of Drosophila proteins that interact with a central nervous system regulatory element. Genes Dev, 1989. 3(8): p. 1130–45.

9. Kudryavtseva, E.I., et al., Identification and characterization of Grainyhead-like epithelial transactivator (GET-1), a novel mammalian Grainyhead-like factor. Dev Dyn, 2003. 226(4): p. 604–17.

10. Ting, S.B., et al., A homolog of Drosophila grainy head is essential for epidermal integrity in mice. Science, 2005. 308(5720): p. 411–3.

11. Wilanowski, T., et al., A highly conserved novel family of mammalian developmental transcription factors related to Drosophila grainyhead. Mech Dev, 2002. 114(1-2): p. 37–50.

12. Prokop, A., et al., Homeotic regulation of segment-specific differences in neuroblast numbers and proliferation in the Drosophila central nervous system. Mech Dev, 1998. 74(1-2): p. 99–110.

13. Uv, A.E., E.J. Harrison, and S.J. Bray, Tissue-specific splicing and functions of the Drosophila transcription factor Grainyhead. Mol Cell Biol, 1997. 17(11): p. 6727–35.

14. Uv, A.E., C.R. Thompson, and S.J. Bray, The Drosophila tissue-specific factor Grainyhead contains novel DNA-binding and dimerization domains which are conserved in the human protein CP2. Mol Cell Biol, 1994. 14(6): p. 4020–31.

15. Chen, W., et al., Grainyhead-like 2 regulates epithelial plasticity and stemness in oral cancer cells. Carcinogenesis, 2016. 37(5): p. 500–10.

16. Mlacki, M., et al., Loss of Grainy head-like 1 is associated with disruption of the epidermal barrier and squamous cell carcinoma of the skin. PLoS One, 2014. 9(2): p. e89247.

17. Xu, H., et al., Clinical implications of GRHL3 protein expression in breast cancer. Tumour Biol, 2014. 35(3): p. 1827–31.

18. Nevil, M., et al., Stable Binding of the Conserved Transcription Factor Grainy Head to its Target Genes Throughout Drosophila melanogaster Development. Genetics, 2017. 205(2): p. 605–620.

19. Jacobs, J., et al., The transcription factor Grainy head primes epithelial enhancers for spatiotemporal activation by displacing nucleosomes. Nat Genet, 2018. 50(7): p. 1011–1020.

20. Nevil, M., et al., Establishment of chromatin accessibility by the conserved transcription factor Grainy head is developmentally regulated. Development, 2020. 147(5).

21. Brody, T. and W.F. Odenwald, Programmed transformations in neuroblast gene expression during Drosophila CNS lineage development. Dev Biol, 2000. 226(1): p. 34–44.

22. Allan, D.W. and S. Thor, Transcriptional selectors, masters, and combinatorial codes: regulatory principles of neural subtype specification. Wiley Interdiscip Rev Dev Biol, 2015. 4(5): p. 505–28.

23. Miyares, R.L. and T. Lee, Temporal control of Drosophila central nervous system development. Curr Opin Neurobiol, 2019. 56: p. 24–32.

24. Doe, C.Q., Temporal Patterning in the Drosophila CNS. Annu Rev Cell Dev Biol, 2017. 33: p. 219–240.

25. Bayraktar, O.A. and C.Q. Doe, Combinatorial temporal patterning in progenitors expands neural diversity. Nature, 2013. 498(7455): p. 449–55.

26. Cenci, C. and A.P. Gould, Drosophila Grainyhead specifies late programmes of neural proliferation by regulating the mitotic activity and Hox-dependent apoptosis of neuroblasts. Development, 2005. 132(17): p. 3835–45.

27. Almeida, M.S. and S.J. Bray, Regulation of post-embryonic neuroblasts by Drosophila Grainyhead. Mech Dev, 2005. 122(12): p. 1282–93.

28. Bello, B.C., F. Hirth, and A.P. Gould, A pulse of the Drosophila Hox protein Abdominal-A schedules the end of neural proliferation via neuroblast apoptosis. Neuron, 2003. 37(2): p. 209–19.

29. Khandelwal, R., et al., Combinatorial action of Grainyhead, Extradenticle and Notch in regulating Hox mediated apoptosis in Drosophila larval CNS. PLoS Genet, 2017. 13(10): p. e1007043.

30. Ghosh, N., et al., Hox gene Abdominal-B uses Doublesex(F) as a cofactor to promote neuroblast apoptosis in Drosophila central nervous system. Development, 2019.

31. Bakshi, A., et al., Sequential activation of Notch and Grainyhead gives apoptotic competence to Abdominal-B expressing larval neuroblasts in Drosophila Central nervous system. PLoS Genet, 2020. 16(8): p. e1008976.

32. Sipani, R. and R. Joshi, Hox genes collaborate with helix-loop-helix factor Grainyhead to promote neuroblast apoptosis along the anterior-posterior axis of the Drosophila larval central nervous system. Genetics, 2022. 222(1).

33. Chai, P.C., et al., Hedgehog signaling acts with the temporal cascade to promote neuroblast cell cycle exit. PLoS Biol, 2013. 11(2): p. e1001494.

34. de Vries, M., et al., Interrogating the Grainyhead-like 2 (Grhl2) genomic locus identifies an enhancer element that regulates palatogenesis in mouse. Dev Biol, 2020. 459(2): p. 194–203.

35. An, H., et al., Pipsqueak family genes dan/danr antagonize nuclear Pros to prevent neural stem cell aging in Drosophila larval brains. Front Mol Neurosci, 2023. 16: p. 1160222.

36. Zacharioudaki, E., et al., Genes implicated in stem cell identity and temporal programme are directly targeted by Notch in neuroblast tumours. Development, 2016. 143(2): p. 219–31.

37. Kuzin, A., et al., Structure and cis-regulatory analysis of a Drosophila grainyhead neuroblast enhancer. Genesis, 2018. 56(3): p. e23094.

38. Negre, N., et al., A comprehensive map of insulator elements for the Drosophila genome. PLoS Genet, 2010. 6(1): p. e1000814.

39. Carleton, J.B., K.C. Berrett, and J. Gertz, Multiplex Enhancer Interference Reveals Collaborative Control of Gene Regulation by Estrogen Receptor alpha-Bound Enhancers. Cell Syst, 2017. 5(4): p. 333–344 e5.

40. Osterwalder, M., et al., Enhancer redundancy provides phenotypic robustness in mammalian development. Nature, 2018. 554(7691): p. 239–243.

41. Berger, C., et al., FACS purification and transcriptome analysis of drosophila neural stem cells reveals a role for Klumpfuss in self-renewal. Cell Rep, 2012. 2(2): p. 407–18.

42. Murphy, P.J. and F. Berger, The chromatin source-sink hypothesis: a shared mode of chromatin-mediated regulations. Development, 2023. 150(21).

43. Korzelius, J., et al., Escargot maintains stemness and suppresses differentiation in Drosophila intestinal stem cells. EMBO J, 2014. 33(24): p. 2967–82.

44. Li, Y., et al., Transcription Factor Antagonism Controls Enteroendocrine Cell Specification from Intestinal Stem Cells. Sci Rep, 2017. 7(1): p. 988.

45. Guo, X., et al., Cell-fate conversion of intestinal cells in adult Drosophila midgut by depleting a single transcription factor. Nat Commun, 2024. 15(1): p. 2656.

46. Neumuller, R.A., et al., Genome-wide analysis of self-renewal in Drosophila neural stem cells by transgenic RNAi. Cell Stem Cell, 2011. 8(5): p. 580–93.

47. Estella, C., D.J. McKay, and R.S. Mann, Molecular integration of wingless, decapentaplegic, and autoregulatory inputs into Distalless during Drosophila leg development. Dev Cell, 2008. 14(1): p. 86–96.

48. Bischof, J., et al., An optimized transgenesis system for Drosophila using germ-line-specific phiC31 integrases. Proc Natl Acad Sci U S A, 2007. 104(9): p. 3312–7.

49. Noro, B., et al., Distinct functions of homeodomain-containing and homeodomain-less isoforms encoded by homothorax. Genes Dev, 2006. 20(12): p. 1636–50.

50. Yang, L., et al., Single molecule fluorescence in situ hybridisation for quantitating post-transcriptional regulation in Drosophila brains. Methods, 2017. 126: p. 166–176.

51. Ramirez, F., et al., High-resolution TADs reveal DNA sequences underlying genome organization in flies. Nat Commun, 2018. 9(1): p. 189.

52. Lopez-Delisle, L., et al., pyGenomeTracks: reproducible plots for multivariate genomic datasets. Bioinformatics, 2021. 37(3): p. 422–423.

