## Supplementary material for "*Drosophila grainyhead* gene and its neural stem cell-specific enhancers show epigenetic synchrony in the cells of the central nervous system": Sup Data_27_02_25.docx

**Supplementary Data**

**Sub-fragmentation of *Int-2BF1***

We split the 1.2 Kb region covered by *Int-2BF1* into two overlapping fragments (Fig S1A, *Int-2BF1A* and *Int-2BF1B*, with a 150bp overlap) and found both of these fragments show NSC-specific expression similar to 1.2 Kb and 4 Kb CREs (Fig S1B-S1C), suggesting that core sequences regulating enhancer activity lie within the overlapping 150bps (*2BF1-AB*) (Fig-S1A). The 1.2 Kb region was then split into three contiguous non-overlapping fragments*, F1A’* (446bp)*, F1B’* (591bp*) and F1AB* (150bp)*,* which were checked for the expression in larval NBs (Table 1). The fragment *F1A’-lacZ* showed a significant expression in NSCs with almost all the cells in the central brain (CB) and a subset of NSCs in the ventral nerve cord (VNC) showed the expression of the reporter (Sup Fig-1B), while *F1B’-lacZ* was much sparse in its expression in CB and had much-reduced expression in VNC (Sup Fig-1C). While 150bp *F1AB*-lacZ was barely visible in the NBs of CB and VNC (Sup Fig-1D). Since *F1AB* was an overlapping region of the *2BF1A* and *2BF1B*, and both had shown strong expression in larval CNS, we considered the possibility that *F1-AB-lacZ was* expressed weakly owing to its small size. Therefore, we trimerised this fragment to generate *2BF1AB-3X-lacZ,* which was found to express in most of the *grh* expressing NSCs of CB and VNC in larval CNS (Fig S1G).

These results collectively suggest that the CNS-specific 4 Kb grh CRE could be narrowed down to its 1.2 Kb subfragment. Its core activity sequences most likely lie within a 150 bp region, and the regions outside 150 bp are also crucial for its overall expression profile.

**Genomic regions screened for *grh enhancers***

We screened 11 non-coding genomic regions bracketed within two modENCODE class I genomic insulator elements identified for *grh* gene [38] (position indicated by red diamonds in Fig 1B).

Insulator-I:Chr 2R: 17800474..17800484

Insulator-II:Chr 2R: 17843080..17843090

The modENCODE stands for Model Organism Encyclopedia of DNA Elements, it was a Consortium effort which mapped functional elements like transcripts, chromatin marks, regulatory factor binding sites, and origins of DNA replication, in two model organisms Drosophila melanogaster and Caenorhabditis elegans.

Grh has a total of 15 introns (Fig 2A). Of these 15, six introns are very small (*int-4*: 66bps; *int-5*: 321bps; *int-6*: 188bps; *int-12*: 93bps; *int-13*: 154bps; *int-15*: 258bps) and therefore were not screened for the potential CREs. The two previously identified NB-specific CREs are *grh-NB* (a 4 Kb CRE also called *Int-2B* here), within this 4 Kb region lies *grh-NRE* (a 1.3 Kb CRE overlapping *Int-2BF1*)*,* and *15B* enhancer (covered by *Int-7*). The remaining 11 genomic regions from 10 introns and 1 non-coding region upstream of exon-1 were screened for enhancer activity (indicated in Fig-2A). Sequences conserved across multiple Drosophila species were amplied out by PCR and fused to beta-galactosidase gene to make the reporter lines. The *Int-10* could not be amplified using PCR, therefore a publically available promoter-GAL4 line covering this region was used to assess the expression pattern of this region in NBs.

Next, we compared the expression of the *enhancer-lacZ* lines with those available for *grh* gene in the Janelia FlyLight resource. There are 14 lines available for *grh* gene in FlyLight portal. Of the 14 lines, only four are available publically through BDSC (BL 45275, BL 71044, BL 64345, BL 69034). We checked all these four lines and observed that except one (BL69034) the expression of all others concurred with our *enhancer-lacZ* lines for their expression in NSCs, when the CNS were counterstained with Dpn.

The remaining ten lines are not publicly available; for nine out of these ten lines, images are only available for adult CNS. Only the GMR40C11 FlyLight portal has images of larval CNS, which shows very few cells expressing in the central brain, which seems to be neurons. This region corresponds to Int-2A, which we screened. We scored for its expression only in NBs and reported that it does not express in NSCs.

It is to be noted that the expressions shown in the FlyLight resource are z-projects of the confocal sections, while what we show in our images are single z-slices, counterstained with Dpn. Moreover, FlyLight portal expression patterns are counterstained with Neurotactin or Neuroglian and do not have any counterstaining with any of the markers for the NSCs or GMC like Dpn or Grh. Therefore, it becomes difficult to directly compare *reporter-lacZ* line expressions with those reported in the FlyLight portal.

Amongst the four lines available, the following are the details.

BL45275 (GMR40A10): This line corresponds to Int-1. We do not see any expression in NBs in this line, nor did Int-1-lacZ show any expression in NSCs.

BL71044 (GMR40G05): corresponds to a region starting from Int-2A and extending into Int-2B (end at our fragment Int-2BF1A). This also overlaps with previously reported *grhNB-lacZ* (shown in Fig-1A)*.* This region shows strong expression in NSCs of CNS, which is the same as what we have reported for *Int-2BF1-lacZ*.

BL64345 (GMR43D01): Since we could not PCR amplify Int-10, we checked the expression of this line and found it to be expressed in Type-II NSCs and a few Type-I NSCs in the thoracic region.

BL69034 (GMR44G11) corresponds to a region starting from Int-11. We see very weak expression in Type-I NSCs and low but consistent expression in Type-II NSCs. We do not see the same in the BL69034 line.

**Details of category three enhancer expression**

The third category included the remaining 2 enhancers, which could not be categorised in the first two categories (***Int-7***, *Int-10*). *Int-7* covered a previously characterised *15B* enhancer, which expressed in all the NSCs but had a variable activity therein, with some NBs showing very high and some showing very low levels of the reporter (Fig S2A and S2F). The *Int-10*, on the other hand, was unique owing to its specific expression, which was mainly confined to all eight Type-II NBs and ten Type-I NSCs in the OLs and approximately eight thoracic NSCs (Fig-S2K-S2L).
