## Supplementary material for "*Drosophila grainyhead* gene and its neural stem cell-specific enhancers show epigenetic synchrony in the cells of the central nervous system": Table II_CREsCoordinates_revised_Feb2024_F.docx

Table No. 2. Collation of the coordinates of the genomic regions tested and their activity (if any), breakpoints for the deletions. New CREs are shown in red, known CREs are highlighted in yellow, sub-fragments of *Int-2BF1* are highlighted in green, and regions which showed no expression in NB or were not screened are shown in black font.

| **Fragment name** | **Release 6 coordinates** | **Size** | **Expression in NBs** |
| --- | --- | --- | --- |
| ***exon1up*** | 17,800,050..17,802,801 | 2752 bp | Expresses |
| ***Int-1*** | 17,803,392..17,808,098 | 4707 bp | No expression |
| ***Int-2A*** | 17,809,655..17,815,304 | 5650 bp | No expression |
| ***Int-2B /grh4*** | 17,815,305..17,819,455 | 4151 bp | Expresses |
| ***grhNB-lacZ*** | 17,815,344..17,819,417 | 4073bp | Expresses |
| ***grh-NRE*** | 17,814,854..17,816,218 | 1364bp | Expresses |
| ***Int-2BF1*** | 17,815,305..17,816,495 | 1191 bp | Expresses |
| ***Int-2BF1A*** | 17,815,305..17,815,904 | 600 bp | Expresses |
| ***Int-2BF1B*** | 17,815,751..17,816,495 | 745bp | Expresses |
| ***Int-2BF1A’*** | 17,815,305..17,815,751 | 446bp | Expresses |
| ***Int-2BF1B’*** | 17,815,904..17,816,495 | 591bp | Expresses |
| ***Int-2BF1AB/F1AB*** | 17,815,751..17,816,904 | 154 bp | Expresses |
| ***Int-2BF2*** | 17,816,496..17,817,995 | 1499bp | No expression |
| ***Int-2BF3*** | 17,817,996..17,819,455 | 1459bp | No expression |
| ***Int-2C*** | 17,819,456..17,820,999 | 1544 bp | Expresses |
| ***Int-3*** | 17,821,412..17,827,133 | 5722 bp | No expression |
| ***Int-4*** | 17,827,329..17,827,394 | 66bp | Not screened |
| ***Int-5*** | 17,828,010..17,828,330 | 321bp | Not screened |
| ***Int-6*** | 17,828,555..17,828,742 | 188bp | Not screened |
| ***Int-7*** | 17,828,906..17,830,996 | 2091bp | Expresses |
| ***grh15B*** | 17,829,129..17,829,979 | 851bp | Expresses |
| ***Int-8*** | 17,831,109..17,831,670 | 562bp | Expresses |
| ***Int-9*** | 17,831,766..17,834,614 | 2849bp | Expresses |
| ***Int-10*** | 17,834,672..17,835,750 | 1079bp | Expresses |
| ***Int-11*** | 17,835,850..17,839,041 | 3192bp | Expresses |
| ***Int-12*** | 17,839,222..17,839,314 | 93bp | Not screened |
| ***Int-13*** | 17,839,440..17,839,593 | 154bp | Not screened |
| ***Int-14*** | 17,839,790..17,840,434 | 645bp | Expresses |
| ***Int-15*** | 17,840,500..17,840,747 | 248bp | Not screened |
| ***CRE Deletions*** | **Breakpoints** | **Size** | **Phenotype** |
| ***Int-2BF1*** | 17,815,158..17,816,534 | 1377bp | Homozygous viable |
| ***Int-2C*** | 17,819,375..17,820,929 | 1554bp | Homozygous viable |
| ***Int-2C’* in double deletion** | 17,819,578..17,820,931 | 1354bp | Homozygous embryonic lethal |
