## Supplementary material for "*Drosophila grainyhead* gene and its neural stem cell-specific enhancers show epigenetic synchrony in the cells of the central nervous system": Table_I_28_02_25.docx

**Table 1: Expression details of new and Known *grh* CREs. CREs from Category I, II and III are shown in green, pink and blue colors**

The first category ncludes the enhancers which show strong and uniform expression in the majority of the NSCs/NBs (*Ex1up,* ***Int-2BF1****, Int-2C, Int-14*) (see images in Fig 1C, 1F-1J, Fig S2E and S2J). The second category CREs were very weak in their expression in NSCs/NBs (*Int-8, Int-9* and *Int-11*) (see images in Fig S2B-S2D and S2G-S2I). The third category included the remaining two enhancers (***Int-7***, *Int-10*), of which *Int-7* showed strong but variable expression in the NBs, and *Int-10* showed strong expression in Type II NSCs/NBs and very limited and weak expression in Type I NBs

| **SNo.** | **New CREs** | **Type I NSCs/NBs (Late L3 stage)** | **Terminal NSCs/NBs in A8-Al0 segments (eL2, mL2, eL3 stages)** | **Type II NSCs/NBs**  **Late L3 stage** | **Wing disc**  **Late L3 stage** |
| --- | --- | --- | --- | --- | --- |
| 1 | *Ex1up* | Strong expression in all NBs  **+ + +** | levels increase exponentially | Expresses | **Broad and strong Expression** |
| 2 | *Int-2C* | Strong expression in NBs of CB and VNC  **+ + + + + +** | levels increase exponentially | Expresses | No expression |
| 3 | *Int-8* | Weak and limited expression in all the NBs  **+** | do not express in the early stages | Expresses | No expression |
| 4 | *Int-9* | Weak and limited expression in all the NBs  **+** | do not express in the early stages | Expresses | No expression |
| 5 | *Int-10* | Weak expression in approximately 8 NBs in the thoracic region (**++**) and strong expression in 10 Type-I NBs of optic lobes (**++**) | do not express in the early stages | **Very strong Expression** | No expression |
| 6 | *Int-11* | Weak and limited expression in CB and VNC NBs  **+** | do not express in the early stages | Expresses | No expression |
| 7 | *Int-14* | Strong expression in all the NBs  **+ + + +** | levels increase exponentially | Expresses | No expression |
|  | **Known CREs** |  |  |  |  |
| 8 | *Int-2BF1* part of *Int-2B/ grh4*/ *grh-NB/ grh-NRE*  Sub-fragment | Strong expression in CB and VNC NBs  **+ + + + + +** | levels increase exponentially | Expresses | No expression |
| 9 | *Int-7 (15B)* | Differential expression in CB and VNC NBs  Some express strongly, some very weakly | levels increase exponentially | Strong Expression | No expression |
