## Supplementary material for "*Drosophila grainyhead* gene and its neural stem cell-specific enhancers show epigenetic synchrony in the cells of the central nervous system": TableIII_ChromatinMarks_enhancers (1).docx

Table III: Detailing the chromatin marks on the *grh* enhancers

| Enhancers | Coordinates | Neural Stem Cells | | | | |
| --- | --- | --- | --- | --- | --- | --- |
|  |  | PolII | Brm | HP1 | Pc | H1 |
| ***Int-7*** | 17,828,906..17,830,996 |  |  |  |  |  |
| ***Int-8*** | 17,831,109..17,831,670 |  |  |  |  |  |
| ***Int-9*** | 17,831,766..17,834,614 |  |  |  |  |  |
| ***Int-10*** | 17,834,672..17,835,750 |  |  |  |  |  |
| ***Int-11*** | 17,835,850..17,839,041 |  |  |  |  |  |
| ***Int-14*** | 17,839,790..17,840,434 |  |  |  |  |  |
| ***exon1up*** | 17,800,050..17,802,801 |  |  |  |  |  |
| Enhancers | Coordinates | Ganglion Mother cells | | | | |
|  |  | PolII | Brm | HP1 | Pc | H1 |
| ***Int-7*** | 17,828,906..17,830,996 |  |  |  |  |  |
| ***Int-8*** | 17,831,109..17,831,670 |  |  |  |  |  |
| ***Int-9*** | 17,831,766..17,834,614 |  |  |  |  |  |
| ***Int-10*** | 17,834,672..17,835,750 |  |  |  |  |  |
| ***Int-11*** | 17,835,850..17,839,041 |  |  |  |  |  |
| ***Int-14*** | 17,839,790..17,840,434 |  |  |  |  |  |
| ***exon1up*** | 17,800,050..17,802,801 |  |  |  |  |  |
| Enhancers | Coordinates | Neurons | | | | |
|  |  | PolII | Brm | HP1 | Pc | H1 |
| ***Int-7*** | 17,828,906..17,830,996 |  |  |  |  |  |
| ***Int-8*** | 17,831,109..17,831,670 |  |  |  |  |  |
| ***Int-9*** | 17,831,766..17,834,614 |  |  |  |  |  |
| ***Int-10*** | 17,834,672..17,835,750 |  |  |  |  |  |
| ***Int-11*** | 17,835,850..17,839,041 |  |  |  |  |  |
| ***Int-14*** | 17,839,790..17,840,434 |  |  |  |  |  |
| ***exon1up*** | 17,800,050..17,802,801 |  |  |  |  |  |
